# Graph abstraction reconciles clustering with trajectory inference through a topology preserving map of single cells

**DOI:** 10.1101/208819

**Authors:** F. Alexander Wolf, Fiona Hamey, Mireya Plass, Jordi Solana, Joakim S. Dahlin, Berthold Göttgens, Nikolaus Rajewsky, Lukas Simon, Fabian J. Theis

**Affiliations:** Helmholtz Center Munich – German Research Center for Environmental Health, Institute of Computational Biology, Neuherberg, Munich, Germany; Department of Haematology and Wellcome Medical Research Council Cambridge Stem Cell Institute, University of Cambridge, Cambridge, United Kingdom; Berlin Institute for Medical Systems Biology, Max-Delbrück Center for Molecular Medicine, Berlin, Germany; Department of Medicine, Karolinska Institutet and Karolinska University Hospital, Stockholm, Sweden; Department of Mathematics, Technische Universität München, Munich, Germany

## Abstract

Single-cell RNA-seq quantifies biological heterogeneity across both discrete cell types and continuous cell transitions. Partition-based graph abstraction (PAGA) provides an interpretable graph-like map of the arising data manifold, based on estimating connectivity of manifold partitions (https://github.com/theislab/paga). PAGA maps provide interpretable discrete and continuous latent coordinates for both disconnected and continuous structure in data, preserve the global topology of data, allow analyzing data at different resolutions and result in much higher computational efficiency of the typical exploratory data analysis workflow — one million cells take on the order of a minute, a speedup of 130 times compared to UMAP. We demonstrate the method by inferring structure-rich cell maps with consistent topology across four hematopoietic datasets, confirm the reconstruction of lineage relations of adult planaria and the zebrafish embryo, benchmark computational performance on a neuronal dataset and detect a biological trajectory in one deep-learning processed image dataset.

## Introduction

Single-cell RNA-seq offers unparalleled opportunities for comprehensive molecular profiling of thousands of individual cells, with expected major impacts across a broad range of biomedical research. The resulting datasets are often discussed using the term transcriptional landscape. However, the algorithmic analysis of cellular heterogeneity and patterns across such landscapes still faces fundamental challenges, for instance, in how to explain cell-to-cell variation. Current computational approaches attempt to achieve this usually in one of two ways [1]. Clustering assumes that data is composed of biologically distinct groups such as discrete cell types or states and labels these with a discrete variable — the cluster index. By contrast, inferring pseudotemporal orderings or trajectories of cells [2–4] assumes that data lie on a connected manifold [5] and labels cells with a continuous variable — the distance along the manifold. While the former approach is the basis for most unsupervised analyses of single-cell data, the latter enables a better interpretation of continuous phenotypes and processes such as development, dose response and disease progression. Here, we unify both viewpoints.

A central example of dissecting heterogeneity in single-cell experiments concerns data that originate from complex cell differentiation processes. However, analyzing such data using pseudotemporal ordering [2, 6–10] faces the problem that biological processes are usually incompletely sampled. As a consequence, experimental data do not conform with a connected manifold and the modeling of data as a continuous tree structure, which is the basis for existing algorithms, has little meaning. This problem exists even in clustering-based algorithms for the inference of tree-like processes [11– 13], which make the generally invalid assumption that clusters conform with a connected tree-like topology. Moreover, they rely on feature-space based inter-cluster distances, like the euclidean distance of cluster means. However, such distance measures quantify biological similarity of cells only at a local scale and are fraught with problems when used for larger-scale objects like clusters. Efforts for addressing the resulting high non-robustness of tree-fitting to distances between clusters [11] by sampling [12, 13] have only had limited success.

Partition-based graph abstraction (PAGA) [14] resolves these fundamental problems by generating graph-like maps of cells that preserve both continuous and disconnected structure in data at multiple resolutions. The data-driven formulation of PAGA allows to robustly reconstruct branching gene expression changes across different datasets and, for the first time, enabled reconstructing the lineage relations of a whole adult animal [15]. Furthermore, we show that PAGA-initialized manifold learning algorithms converge faster, produce embeddings that are more faithful to the global topology of high-dimensional data and introduce an entropy-based measure for quantifying such faithfulness. Finally, we show how PAGA abstracts transition graphs, for instance, from RNA velocity and compare to previous trajectory-inference algorithms.

## Results

### PAGA maps discrete disconnected and continuous connected cell-to-cell variation

Both established manifold learning techniques and single-cell data analysis techniques represent data as a neighborhood graph of single cells *G* = (*V, E*), where each node in *V* corresponds to a cell and each edge in *E* represents a neighborhood relation (Figure 1) [3, 16–18]. However, the complexity of *G* and noise-related spurious edges make it both hard to trace a putative biological process from progenitor cells to different fates and to decide whether groups of cells are in fact connected or disconnected. Moreover, tracing isolated paths of single cells to make statements about a biological process comes with too little statistical power to achieve an acceptable confidence level. Gaining power by averaging over distributions of single-cell paths is hampered by the difficulty of fitting realistic models for the distribution of these paths.

**Figure 1.**
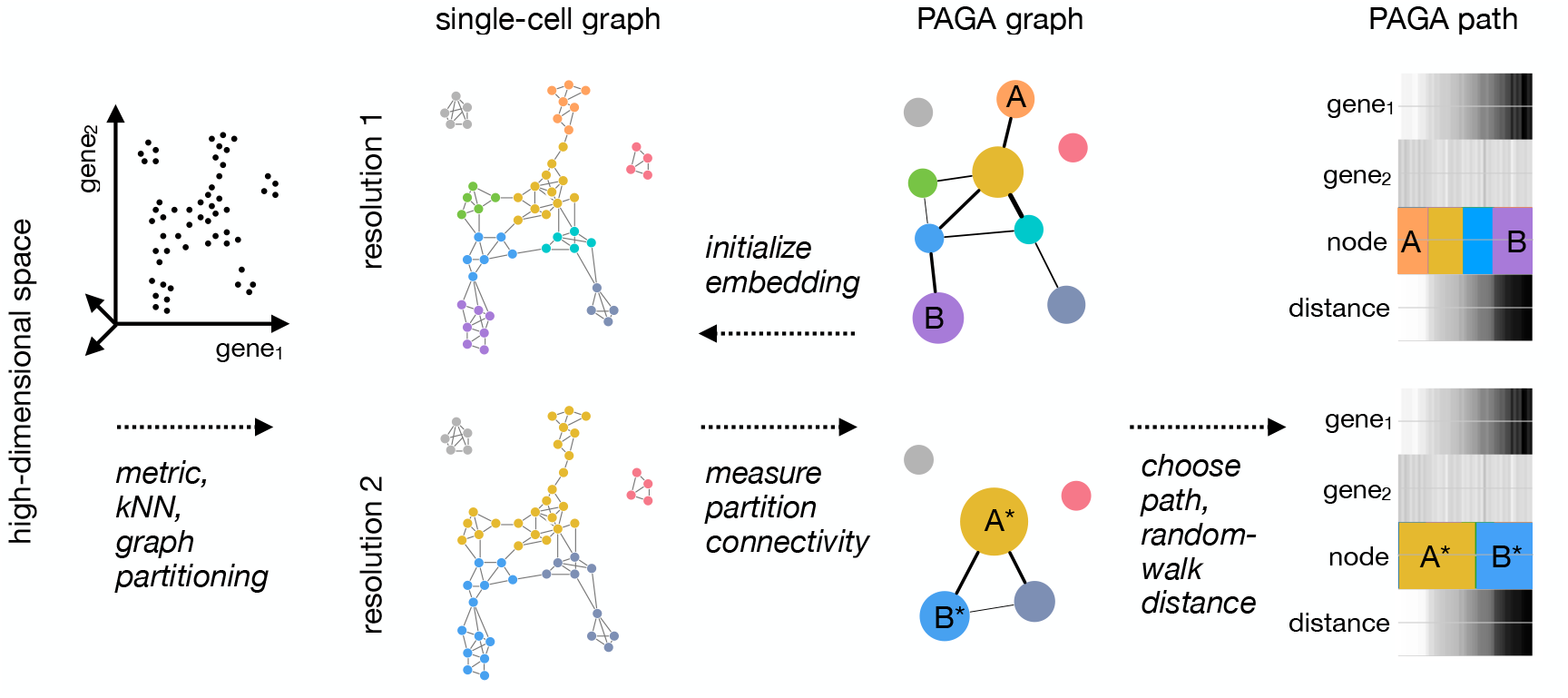
Partition-based graph abstraction generates a topology-preserving map of single cells. High-dimensional gene expression data is represented as a kNN graph by choosing a suitable low-dimensional representation and an associated distance metric for computing neighborhood relations — in most of the paper we use PCA-based representations and Euclidean distance. The kNN graph is partitioned at a desired resolution where partitions represent groups of connected cells. For this, we usually use the Louvain algorithm, however, partitions can be obtained in any other way, too. A PAGA graph is obtained by associating a node with each partition and connecting each node by weighted edges that represent a statistical measure of connectivity between partitions, which we introduce in the present paper. By discarding spurious edges with low weights, PAGA graphs reveal the denoised topology of the data at a chosen resolution and reveal its connected and disconnected regions. Combining high-confidence paths in the PAGA graph with a random-walk based distance measure on the single-cell graph, we order cells within each partition according to their distance from a root cell. A PAGA path then averages all single-cell paths that pass through the corresponding groups of cells. This allows to trace gene expression changes along complex trajectories at single-cell resolution.

We address these problems by developing a statistical model for the connectivity of groups of cells, which we typically determine through graph-partitioning [18–20] or alternatively through clustering or experimental annotation. This allows us to generate a simpler PAGA graph *G^*^* (Figure 1) whose nodes correspond to cell groups and whose edge weights quantify the connectivity between groups. The statistical model considers groups as connected if their number of inter-edges exceeds a fraction of the number of inter-edges expected under random assignment. The connection strength can be interpreted as confidence in the presence of an actual connection and allows discarding spurious, noise-related connections (Supplemental Note 1). While *G* represents the connectivity structure of data at single-cell resolution, the PAGA graph *G^*^* represents the connectivity structure of data at the chosen coarser resolution of the partitioning and allows to identify connected and disconnected regions of data. Following paths along nodes in *G^*^* means following an ensemble of single-cell paths that pass through the corresponding cell groups in *G*. By averaging over such an ensemble of single-cell paths, it becomes possible to trace a putative biological process from a progenitor to fates in a way that is robust to spurious edges, provides statistical power and is consistent with basic assumptions on a biological trajectory of cells (Supplemental Note 2). Note that by varying the resolution of the partitioning, PAGA generates PAGA graphs at multiple resolutions, which enables a hierarchical exploration of data (Figure 1, Supplemental Note 1.3).

To trace gene dynamics at single-cell resolution, we extended existing random-walk based distance measures (Supplemental Note 2, Reference [8]) to the realistic case that accounts for disconnected graphs. By following high-confidence paths in the abstracted graph *G^*^* and ordering cells within each group in the path according to their distance *d* from a progenitor cell, we trace gene changes at single-cell resolution (Figure 1). Hence, PAGA covers both aspects of clustering and pseudotemporal ordering by providing a coordinate system (*G^*^, d*) that allows us to explore variation in data while preserving its topology (Supplemental Note 1.6). PAGA can thus be viewed as an easily-interpretable and robust way of performing topological data analysis [10, 21] (Supplemental Note 3).

### PAGA-initialized manifold learning produces topology-preserving single-cell embeddings

The computationally almost cost-free coarse-resolution embeddings of PAGA can be used to initialize established manifold learning and graph drawing algorithms like UMAP [22] and ForceAtlas2 (FA) [23]. This strategy is used to generate the single-cell embeddings throughout this paper. In contrast to the results of previous algorithms, PAGA-initialized single-cell embeddings are faithful to the global topology, which greatly improves their interpretability. To quantify this claim, we took a classification perspective on embedding algorithms and developed a cost function KL_geo_ (Box and Supplemental Note 4), which captures faithfulness to global topology by incorporating geodesic distance along the representations of data manifolds in both the high-dimensional and the embedding space, respectively. Independent of this, PAGA-initialized manifold learning converges about 6 times faster with respect to established cost functions in manifold learning (Supplemental Figure 10).

**Box** Taking a classification view on embedding algorithms, we quantify how faithful an embedding is to the global topology of the high-dimensional data by comparing the distributions *P* and *Q* of edges in the high-dimensional and embedding spaces using a weighted Kullback-Leibler divergence

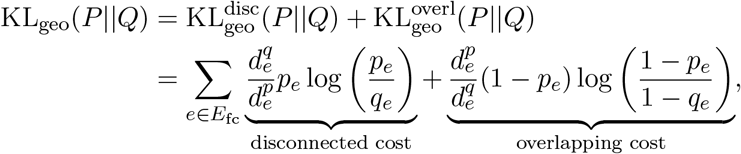

where *p_e_* and *q_e_* are the probabilities for an edge being present in the kNN graphs in the high-dimensional and embedding spaces, respectively. Analogously, 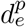 and 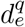 denote random-walk based estimators of geodesic distances on the manifolds in these spaces, respectively. *E*_fc_ denotes the edge set of the fully connected graph (Supplemental Note 4, Supplemental Figure 10).

### PAGA consistently predicts developmental trajectories and gene expression changes in datasets related to hematopoiesis

Hematopoiesis represents one of the most extensively characterised systems involving stem cell differentiation towards multiple cell fates and hence provides an ideal scenario for applying PAGA to complex manifolds. We applied PAGA to simulated data (Supplemental Note 5) for this system and three experimental datasets: 2,730 cells measured using MARS-seq [24], 1,654 cells measured using Smart-seq2 [25] and 44,802 cells from a 10x Genomics protocol [26]. These data cover the differentiation from stem cells towards, cell fates including erythrocytes, megakaryocytes, neutrophils, monocytes, basophils and lymphocytes.

The PAGA graphs (Figure 2) capture known features of hematopoiesis, such as the proximity of megakaryocyte and erythroid progenitors and strong connections between monocyte and neutrophil progenitors. Under debate is the origin of basophils. Studies have suggested both that basophils originate from a basophil-neutrophil-monocyte progenitor or, more recently, from a shared erythroid-megakaryocyte-basophil progenitor [27, 28]. The PAGA graphs of the three experimental datasets highlight this ambiguity. While the dataset of Paul *et al.* falls in the former category, Nestorowa *et al.* falls in the latter and Dahlin *et al.*, which has by far the highest cell numbers and the densest sampling, allows us to see both trajectories. Aside from this ambiguity that can be explained by insufficient sampling in Paul *et al.* and Nestorowa *et al.*, even with the very different experimental protocols and vastly different cell numbers the PAGA graphs show consistent topology between the three datasets. Beyond consistent topology between cell subgroups, we find consistent continuous gene expression changes across all datasets — we observe changes of erythroid maturity marker genes (*Gata2*, *Gata1*, *Klf1*, *Epor* and *Hba-a2*) along the erythroid trajectory through the PAGA graphs and observe sequential activation of these genes in agreement with known behaviour. Activation of neutrophil markers (*Elane*, *Cepbe* and *Gfi1*) and monocyte markers (*Irf8*, *Csf1r* and *Ctsg*) are seen towards the end of the neutrophil and monocyte trajectories, respectively. While PAGA is able to capture the dynamic transcriptional processes underlying multi-lineage hematopoietic differentiation, previous algorithms fail to produce robust or meaningful results (Supplemental Figures 8 and 9).

**Figure 2.**
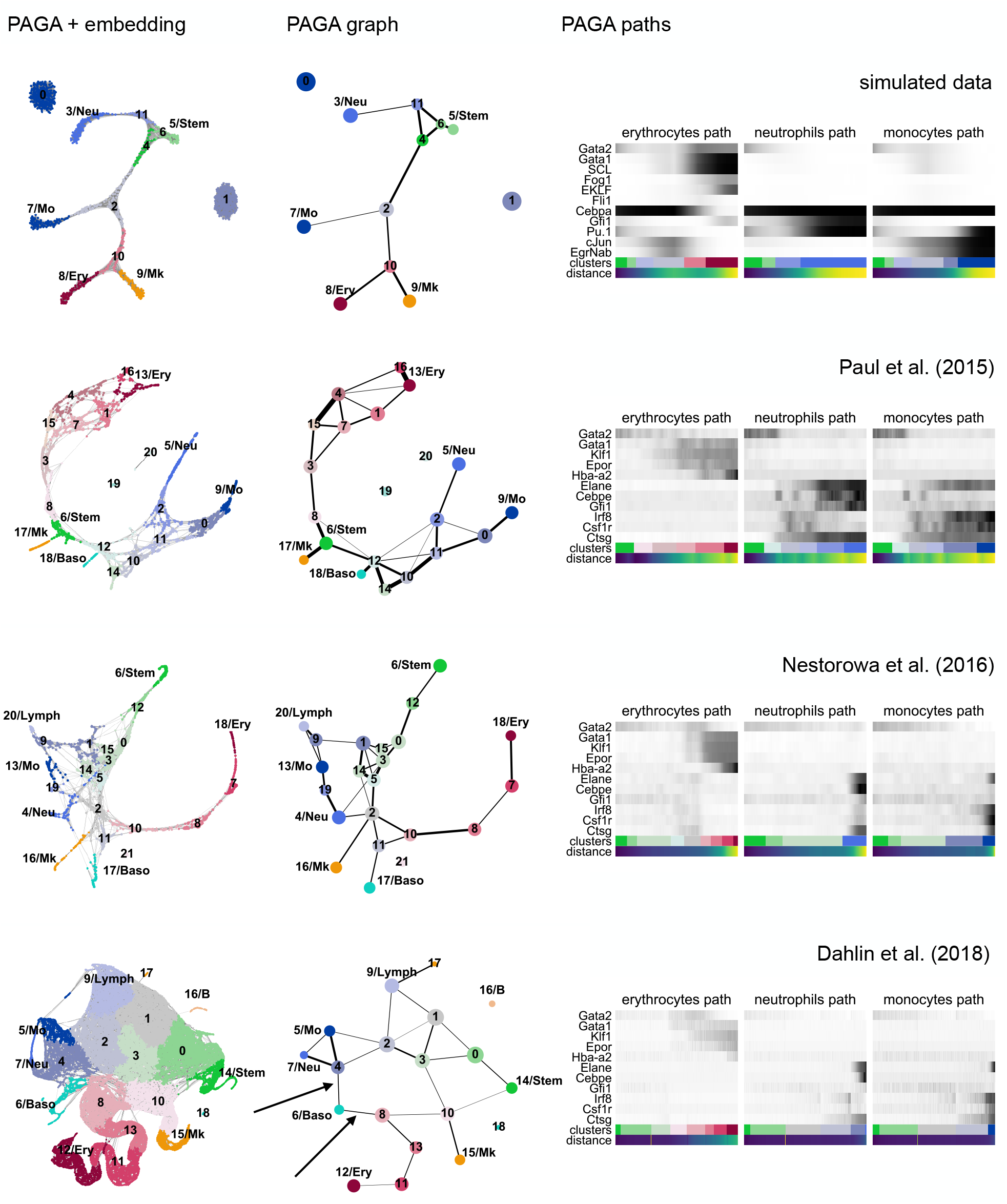
PAGA consistently predicts developmental trajectories and gene expression changes across datasets for hematopoiesis. The three columns correspond to PAGA-initialized single-cell embeddings, PAGA graphs and gene changes along PAGA paths. The four rows of panels correspond to simulated data (Supplemental Note 5) and data from Paul *et al.* [24], Nestorowa *et al.* [25] and Dahlin *et al.* [26], respectively. The arrows in the last row mark the two trajectories to Basophils. One observes both consistent topology of PAGA graphs and consistent gene expression changes along PAGA paths for 5 erythroid, 3 neutrophil and 3 monocyte marker genes across all datasets. The cell type abbreviations are: Stem for stem cells, Ery for erythrocytes, Mk for megakaryocytes, Neu for neutrophils, Mo for monocytes, Baso for basophils, B for B cells, Lymph for lymphocytes.

### PAGA maps single-cell data of whole animals at multiple resolutions

Recently, Plass *et al.* [15] reconstructed the first cellular lineage tree of a whole adult animal, the flatworm *Schmidtea mediterranea*, using PAGA on scRNA-seq data from 21,612 cells. While Plass *et al.* focussed on the tree-like subgraph that maximizes overall connectivity — the minimum spanning tree of *G^*^* weighted by inverse PAGA connectivity — here, we show how PAGA can be used to generate maps of data at multiple resolutions (Figure 3a). Each map preserves the topology of data, in contrast to state-of-the-art manifold learning where connected tissue types appear as either disconnected or overlapping (Figure 3b). PAGA’s multi-resolution capabilities directly addresses the typical practice of exploratory data analysis, in particular for single-cell data: data is typically reclustered in certain regions where a higher level of detail is required.

**Figure 3.**
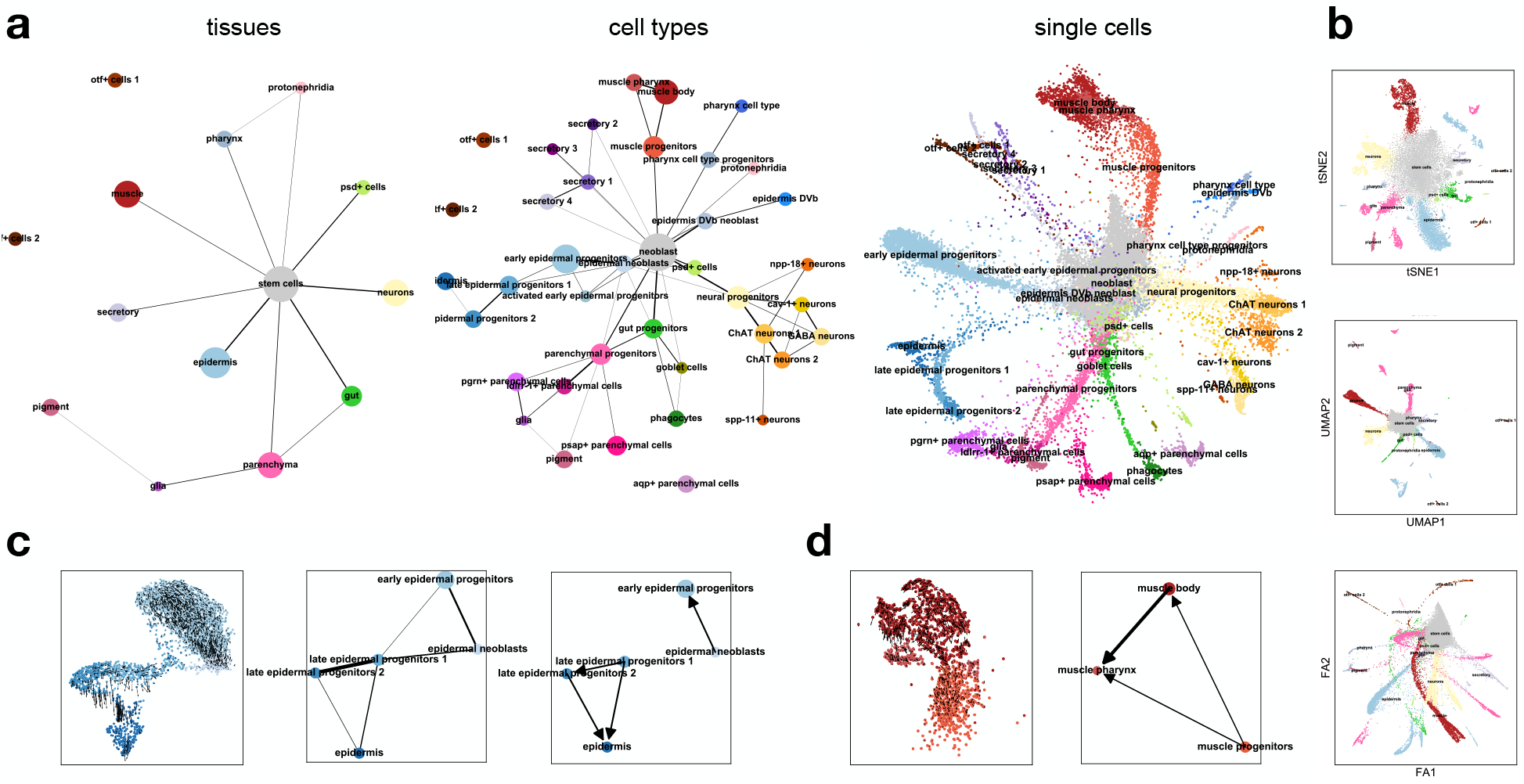
PAGA applied to a whole adult animal. **a,** PAGA graphs for data for the flatworm *Schmidtea mediterranea* [15] at tissue, cell type and single-cell resolution. Only by initializing a single-cell embedding with the embedding of the cell-type PAGA graph, it is possible to obtain a topologically meaningful embedding. Note that the PAGA graph is the same as in Reference [15], only that here, we neither highlight a tree subgraph nor used the corresponding tree layout for visualization. **b,** Established manifold learning for the same data. **c, d,** Predictions of RNA velocity evaluated with PAGA for two example lineages: epidermis and muscle. We show the RNA velocity arrows plotted on a single-cell embedding, the standard PAGA graph representing the topological information (only epidermis) and the PAGA graph representing the RNA velocity information.

### PAGA abstracts information from RNA velocity

Even through the connections in PAGA graphs often correspond to actual biological trajectories, this is not always the case. This is a consequence of PAGA being applied to kNN graphs, which solely contain information about the topology of data. Recently, it has been suggested to also consider directed graphs that store information about cellular transition based on RNA velocity [29]. To include this additional information, which can add further evidence for actual biological transitions, we extend the undirected PAGA connectivity measure to such directed graphs (Supplemental Note 1.2) and use it to orient edges in PAGA graphs (Figure 3c). Due the relatively sparsely sampled, high-dimensional feature space of scRNA-seq data, both fitting and interpreting an RNA velocity vector without including information about topology — connectivity of neighborhoods — is practically impossible. PAGA provides a natural way of abstracting both topological information and information about RNA velocity.

Next, we applied PAGA to 53,181 cells collected at different developmental time points (embryo days) from the zebrafish embryo [30]. The PAGA graph for partitions corresponding to embryo days accurately recovers the chain topology of temporal progression, whereas the PAGA graph for cell types provide easily interpretable overviews of the lineage relations (Figure 4a). Initializing a ForceAtlas2 layout with PAGA coordinates from fine cell types automatically produced a corresponding, interpretable single-cell embedding (Figure 4a). Wagner *et al.* [30] both applied an independently developed computational approach with similarities to PAGA (Supplemental Notes 3) to produce a coarse-grained graph and experimentally validated inferred lineage relations. Comparing the PAGA graph for the fine cell types to the coarse-grained graph of Wagner *et al.* reproduced their result with high accuracy (Figure 4b).

**Figure 4.**
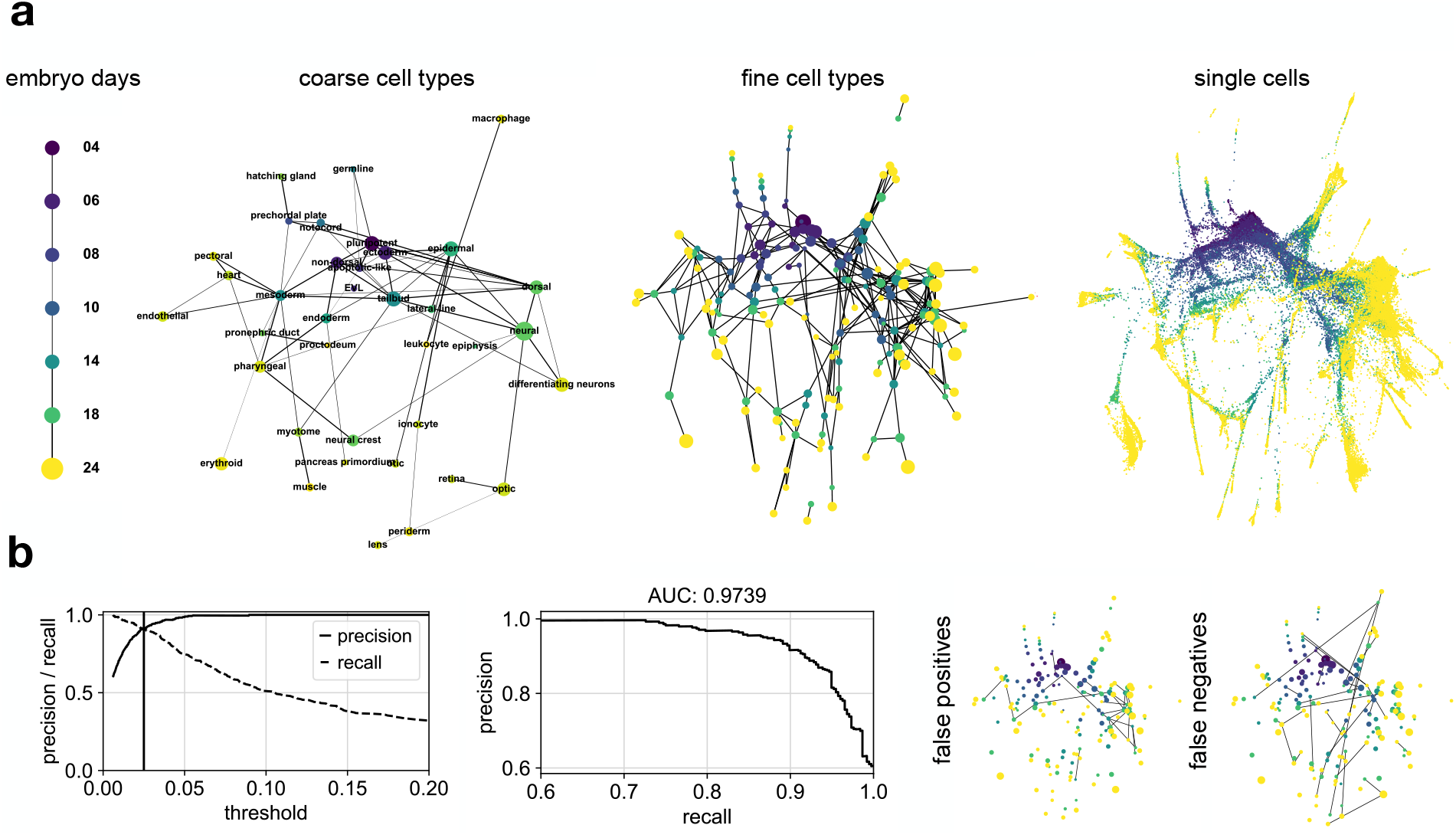
PAGA applied to zebrafish embryo data of Wagner *et al.* [30]. **a,** PAGA graphs obtained after running PAGA on partitions corresponding to embryo days and coarse cell types and a PAGA-initialized single-cell embedding colored by the same quantities. **b,** Performance measurements of the PAGA prediction compared to the reference graph of Wagner *et al.* show high accuracy. False positive edges and false negative edges for the threshold indicate by a vertical line in the left panel are also shown.

### PAGA increases computational efficiency and interpretability in general exploratory data analysis and manifold learning

Comparing the runtimes of PAGA with the state-of-the-art UMAP [22] for 1.3 million neuronal cells of 10x Genomics [31] we find a speedup of about 130, which enables interactive analysis of very large-scale data (90 s versus 191 min on 3 cores of a small server, tSNE takes about 12 10 h). For complex and large data, the PAGA graph generally provides a more easily interpretable visualization of the clustering step in exploratory data analysis, where the limitations of two-dimensional representations become apparent (Supplemental Figure 12). PAGA graph visualizations can be colored by gene expression and covariates from annotation (Supplemental Figure 13) just as any conventional embedding method.

### PAGA is robust and qualitatively outperforms previous lineage reconstruction algorithms

To assess how robustly graph and tree-inference algorithms recover a given topology, we developed a measure for comparing the topologies of two graphs by comparing the sets of possible paths on them (Supplemental Note 1.3 and 1.4, Supplemental Figure 4). Sampling widely varying parameters, which leads to widely varying clusterings, we find that the inferred abstraction of topology of data within the PAGA graph is much more robust than the underlying graph clustering algorithm (Supplemental Figure 5). While graph clustering alone is, as any clustering method, an ill-posed problem in the sense that many highly degenerate quasi-optimal clusterings exist and some knowledge about the scale of clusters is required, PAGA is not affected by this.

Several algorithms [6, 11–13] have been proposed for reconstructing lineage trees (Supplemental Note 3, [4]). The main caveat of these algorithms is that they, unlike PAGA, try to explain any variation in the data with a tree-like topology. In particular, any disconnected distribution of clusters is interpreted as originating from a tree. This produces qualitatively wrong results already for simple simulated data (Supplementary Figure 6) and only works well for data that clearly conforms with a tree-like manifold (Supplementary Figure 7). To establish a fair comparison on real data with the recent popular algorithm, Monocle 2, we reinvestigated the main example of Qiu *et al.* [6] for a complex differentiation tree. This example is based on the data of Paul *et al.* [24] (Figure 2), but with cluster 19 removed. While PAGA identifies the cluster as disconnected with a result that is unaffected by its presence, the prediction of Monocle 2 changes qualitatively if the cluster is taken into account (Supplementary Figure 8). The example illustrates the general point that real data almost always consists of dense and sparse — connected and disconnected — regions, some tree-like, some with more complex topology.

## Discussion

In view of an increasing number of large datasets and analyses for even larger merged datasets, PAGA fundamentally addresses the need for scalable and interpretable maps of high dimensional data. In the context of the Human Cell Atlas [32] and comparable databases, methods for their hierarchical, multi-resolution exploration will be pivotal in order to provide interpretable accessibility to users. PAGA allows for the first time to present information about clusters or cell types in an unbiased, data-driven coordinate system by representing these in PAGA graphs. In the context of the recent advances of the study of simple biological processes that involve a single branching [7, 8], PAGA provides a similarly robust framework for arbitrarily complex topologies. In view of the fundamental challenges of single-cell resolution studies due to technical noise, transcriptional stochasticity and computational burden, PAGA provides a general framework for extending studies of the relations among single cells to relations among noise-reduced and computationally tractable groups of cells. This could facilitate obtaining clearer pictures of underlying biology.

In closing, we note that PAGA not only works for scRNA-seq based on distance metrics that arise from a sequence of chosen preprocessing steps, but can also be applied to any learned distance metric. To illustrate this point, we used PAGA for single-cell imaging data when applied on the basis of a deep-learning based distance metric. Eulenberg *et al.* [33] showed that a deep learning model can generate a feature space in which distances reflect the continuous progression of cell cycle. Using this, PAGA correctly identifies the biological trajectory through the interphases of cell cycle while ignoring a cluster of damaged and dead cells (Supplemental Note 5.6).

## Code and Data availability

PAGA as well as all processing steps used within the analyses are available within Scanpy [34]: https://github.com/theislab/scanpy. The analyses and results of the present paper are available from https://github.com/theislab/paga. Data is linked from https://github.com/theislab/paga.

## Acknowledgements

We thank N. Yosef and D. Wagner for stimulating discussions, S. Tritschler for valuable feedback when testing the code and M. Luecken for comments on graph partitioning algorithms. F.A.W. acknowledges support by the Helmholtz Postdoc Programme, Initiative and Networking Fund of the Helmholtz Association. J.S.D. is supported by a grant from the Swedish Research Council. Work in B.G.’s laboratory is supported by grants from Wellcome, Bloodwise, Cancer Research UK, NIH-NIDDK, and core support grants by Wellcome to the Cambridge Institute for Medical Research and Wellcome-MRC Cambridge Stem Cell Institute. F.K.H. is the recipient of a Medical Research Council PhD Studentship. The work from M.P., J.S., and N.R. was funded by the German Center for Cardiovascular Research (DZHK BER 1.2 VD) and the DFG (grant RA 838/5-1). F.J.T. is supported by the German Research Foundation (DFG) within the Collaborative Research Centre 1243, Subproject A17.

## Supplemental Notes

### Supplemental Note 1: Theoretical background of PAGA

The simplest measure for connectivity of two partitions of *G* is the number of connecting edges between two partitions. However, this number alone depends strongly on the partition sizes, which prevents its meaningful interpretation. Instead, we compute a statistic of this number that measures confidence in an actual connection between two partitions of *G*, as opposed to a connection that is based on spurious edges. Hence, in the visualization of PAGA graphs, edge width should be interpreted as a measure of connectivity whose strength indicates the presence of an actual connection.

Note that topological data analysis (TDA) [21] uses clustering algorithms that lead to overlapping clusters and by that circumvents a statistical definition of a connectivity measure: two clusters are connected if they have finite overlap. In contrast to the easily interpretable, essentially parameter-free and computational efficient Louvain algorithm for modularity optimization [19] — which therefore has become the standard for single-cell analysis [18] — TDA comes with problems in all three respects and is hence not widely used despite it’s recent proposition for scRNA-seq [10]. Also, to date, overlapping graph partitioning algorithms, which also provide a notion of connectivity based on overlaps are, to date, no practical alternative.

#### Supplemental Note 1.1: PAGA for mapping connectivity between partitions

In order to derive a statistical model for connectivity between partitions of kNN graphs of data we need to study the distribution of inter-edges between partitions of the graph. In the simplest case, the model is a distribution conditioned on partition sizes. However, a priori, it is somewhat unclear on which degree distribution of a graph a model for inter-edges should be based. By construction, kNN graphs display a degree distribution “close to a constant degree *k*”. However, this is not an exact constraint and can be violated. As model systems, we investigate graphs with constant and arbitrary degree distribution. We will then choose the appropriate model by comparing model predictions with the empirical estimates of inter-edge distributions that we obtain from kNN graphs.

##### Graphs with constant degree distribution

Consider the case of a partitioned directed graph *G* = (*V, E*) with *h_i_* half-edges or “edge stubs” attached to nodes in a given partition *i*, *h_j_* half-edges attached to nodes in a partition *j* and a total of 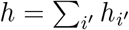 half-edges, which we require to be even. Connected among each other, the half-edges give rise to the 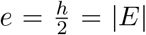 edges of *G*. Assuming half-edges to be distinguishable and randomly connecting half-edges constrained to a constant outdegree distribution of *k* = 1 in *i* and indegree distribution of *k* = 1 in *j*, one has 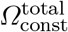 possibilities of combining edges and 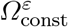 possibilities so that exactly *ε_ij_* inter-edges from partitions *i* and *j* are obtained, with

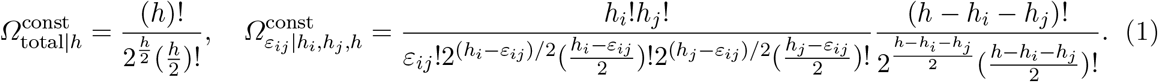

The resulting probability is 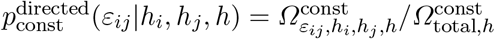. ^1^ We now wish an estimate for the number of inter-edges that is useful also for undirected graphs, hence, the sampling procedure should be symmetric between *i* and *j* and the distribution of interest is the one of the summed inter-edges *ε* = *ε_ij_* + *ε_ji_* when connecting outgoing edges both from *i* and *j*,

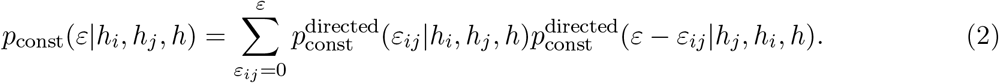

##### Graphs with arbitrary degree distribution

Consider again a partitioned directed graph *G* = (*V, E*), but now with *e* = |*E*| edges and *n* = |*V*| nodes. Imagine we have *e_i_* dangling outgoing edges attached to *n_i_* nodes in partition *i* and we randomly connect each of the dangling edges to a random node in the graph. Enumeration as before gives 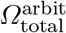 possibilities of connecting these edges and 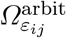 possibilities so that exactly *ε_ij_* inter-edges from partition *i* to *j* are obtained,

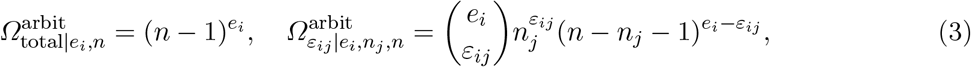

where 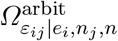 is the product of the possibilities for *ε_ij_* inter-edges from *i* to *j* and the total possibilities of edges among the remaining nodes *i*. Upon definition of 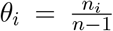, the resulting probability becomes a binomial

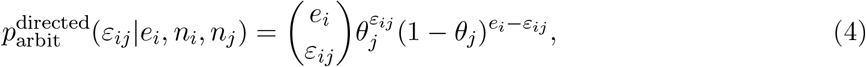

or equivalently,

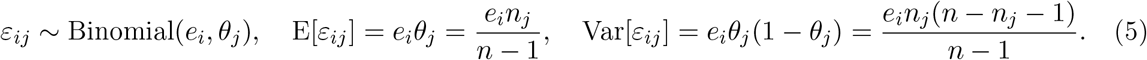

**Supplemental Figure 1.**
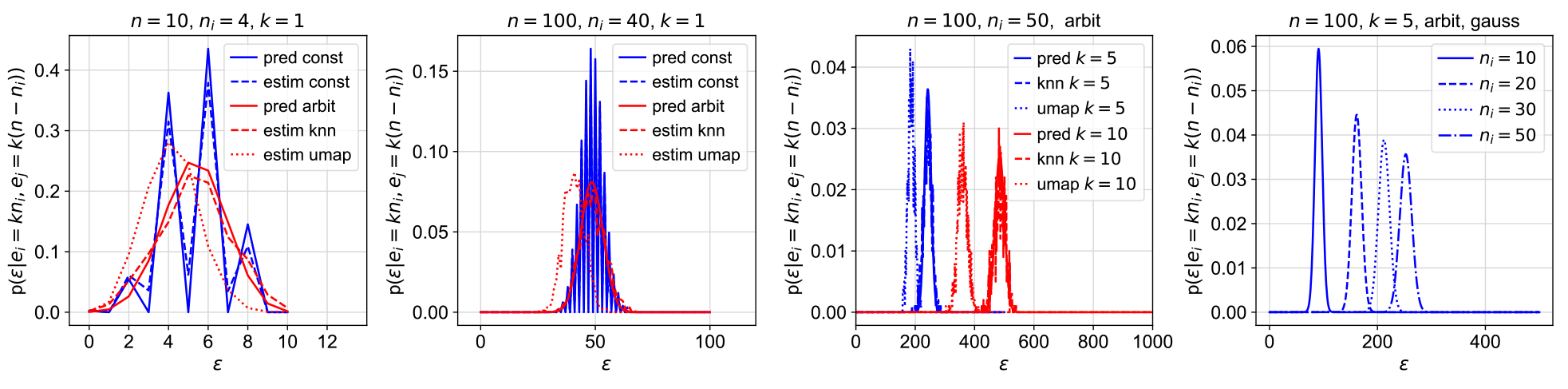
Distribution of inter edges for random bipartitions of fixed degree and for knn graphs. Shown are both sampling basted estimates and model predictions for different parameters.

We can interpret *ε_ij_* as the number of “hitting” partition *j* when randomly distributing the *e_i_* edges of partition *i* where *θ_j_* is the probability for “hitting” *j* for a single edge from *i*.

As before, we wish the sampling procedure to be symmetric between *i* and *j*, hence are interested in the distribution of *ε* = *ε_ij_* + *ε_ji_*, which reads

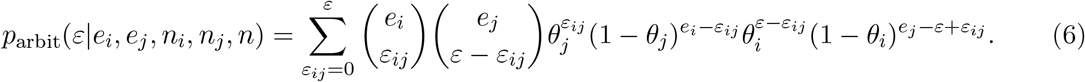

Often, we consider sufficiently large partitions and the binomial distributions become well-approximated by Gaussians with means and variances as in (5). Hence, *ε* is well-approximated as the sum of two Gaussian random variables

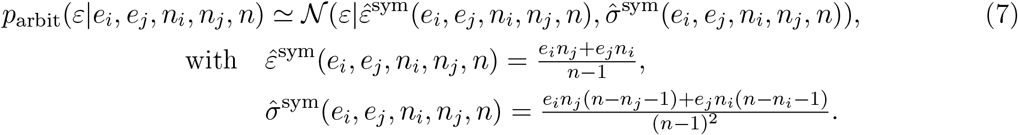

Assuming a knn graph with, at least on average, *e_i_* = *kn_i_* and *e_j_* = *kn_j_*, this simplifies further

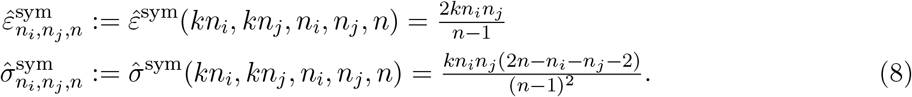

##### Comparing model predictions with sampling based estimates for kNN graphs

To assess how well models (2) and (7) capture the distribution of the number of inter-edges *ε* in sampling-based simulations, we studied three sampling-based models for bipartitioned graphs. The first model randomly connects half-edges in a bipartitioned set of nodes to simulate constant-outdegree *k* = 1 graphs with partition sizes *n_i_* and *n_j_* = *n – n_i_*. The second and third model fit kNN graphs to data sampled from a Gaussian and proceed with randomly partitioning this graph by assigning nodes to random binary partition label — again for fixed *n_i_* and *n_j_* = *n – n_i_*. The third model is equivalent to the second but uses instead of the non-symmetric kNN graph, the symmetrized kNN graph — as results, for instance, in UMAP from the fuzzy union of local simplicial sets to each data point [22].

Results of the sampling simulations based on estimates of 1000 samples show the following findings (Supplemental Figure 1).

1. There is a strong difference in the functional form of distributions for constant and arbitrary degree graphs for small numbers of nodes (*n* = 10 *n_i_* = 4). For higher values (*n* = 100, *n_i_* = 40), both distributions approach Gaussians (two left panels of Supplemental Figure 1).
2. The constant-degree sampling based estimate agrees well with the prediction for the constant-degree model (2), the kNN-fitting based sampling estimate agrees well with the prediction of the arbitrary-degree model (7) (two left panels of Supplemental Figure 1).
3. If one evaluates the kNN graph as in UMAP, one obtains a an empirical distribution that is no longer well-described by (7) (center panels of Supplemental Figure 1).
4. The number of inter-edges depends on the partition-sizes (right panel of Supplemental Figure 1). The relation on *n_i_* and *n_j_* = *n − n_i_* can be seen to be quadratic from (2).

As mentioned, kNN graphs do not have constant-degree *k* distributions as nearest-neighbor-relations among two observations 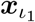 and 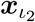 are often not symmetric. Still kNN graphs are somewhat “close” to a constant degree *k* distribution. Supplemental Figure 1 provides strong evidence that this is also the case when using model (7), which, in principle, accounts for arbitrary degree distributions. This can also be theoretically understood by acknowledging that also in the model, strong deviations from degree *k* are unlikely, in particular for sparse graphs with *k* ≪ *n* which implies a comparatively low variance of *k*.

##### Statistical test for disconnectedness and PAGA connectivity measure

Given equation (7), it is straight-forward to write down a hypothesis test for disconnectedness of two partitions *i* and *j* with the null hypothesis that edges of *i* and *j* are randomly connected among each other. One can reject the hypothesis of connectedness at a p-value *p* with an observed inter-edge number 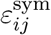, observed edge numbers of partitions *e_i_*, *e_j_* and partition sizes *n_i_*, *n_j_*

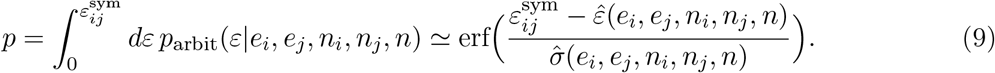

In order to define a connectivity measure for a PAGA graph, this p-value has the desired property of varying in [0, 1] taking large values if it is likely that partitions are connected and taking small values if it is unlikely. Given that we expect the null model of random connections to strongly overestimate inter-edge numbers when applied in practice, the exponential variation of the p-value hampers interpretation and visualization of PAGA graphs, in which edge thickness should indicate connectivity. Using the p-values logarithmized version resolves the exponential variation, but does not vary in [0, 1] anymore.

So, instead of using the p-value for quantifying connectivity *c* of partitions in a PAGA graph, we suggest a linear function of the test statistic

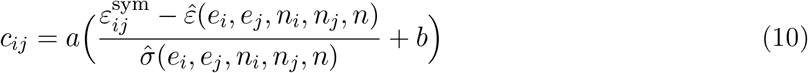

such that the connectivity measure takes values in [0, 1] with *ε* = 0 corresponding to connectivity *c* = 0 and *ε ≥ ε*̂ corresponding to connectivity *c* = 1. Solving these conditions for *a* and *b* results in

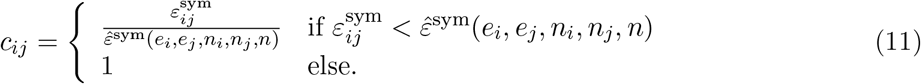

This provides the basic model for “PAGA connectivity” through this paper — below we discuss how this measure could presumably be improved.

##### Relation to modularity

In practice we are not interested in random partitions but in partitions that show stronger intra-partition than inter-partition connectivity. Typically, one uses modularity optimization [19, 35, 36] to compute such a partitioning. To conform with our previous notation, we not only give modularity *m_ij_* but also a symmetrized version *m*_ij_^sym^ of it

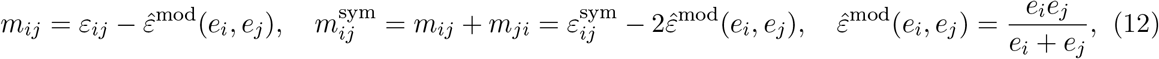

where *e_i_* is the number of outgoing edges of nodes in partition *i*, *ε_ij_* is the number of edges from partition *i* to partition *j* and *ε_ij_* = *ε_ij_* + *ε_ji_* is the number of inter-edges, as before. Note that in modularity optimization algorithms, the modularity measure is only evaluated for the same partition *i* = *j*. Here, *ε*̂ ^mod^(*e_i_, e_j_*) is the expected number of inter-edges between partitions when randomly connecting edges to edges — and not edges to nodes as in (7). This results in a probability 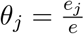 for connecting a given edge in partition *i* to an edge in partition *j*, hence *e_i_θ_j_* expected edges from *i* to *j*. If one assumes a directed constant-outdegree *k* kNN graph with *e_i_* = *kn_i_*, *ε*̂ ^sym^(*e_i_, e_j_*) and 2*ε*̂ ^mod^(*e_i_, e_j_*) agree up to a small difference in the denominator — (*e_i_* + *e_j_ – k*) versus (*e_i_* + *e_j_*) — which comes from avoiding self-loops in *ε*̂ ^sym^(*e_i_, e_j_*) — a property of kNN graphs fitted to data.

##### Feature-space based connectivity

Let us now investigate whether we can relate the graph-based statistical measures of connectivity to a notion of connectivity for the feature space *χ* of observations ***x***_*ι*_. This will also provide the basis for systematically benchmarking the previous graph-based measures on simulated data.

Consider the Gaussian mixture

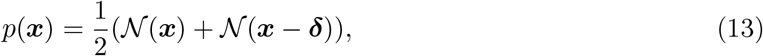

where 𝒩 (***x***) is the standard normal distribution (***μ***_1_ = **0**, ***Σ*** = diag(**1**)) and ***μ***_2_ = ***δ*** denotes the mean of a second shifted normal distribution. Data sampled from this model show two clusters if the distance between the cluster centers *δ* = |***δ*|** is large enough. Otherwise, upon visual inspection, the data “appear to be connected” even though model selection on Gaussian mixture models might still select (13) as the model that most likely explains the data. While the structure of the model (13) should be considered topologically disconnected as it does not rely on the parametrization of a connected manifold, clearly, a kNN-graph fitted to data sampled is strongly connected across clusters if the cluster centers are close enough.

Can we define a notion of connectivity on the level of the model that reflects the connectivity observed in the knn graph? We suggest to define a “connected region” of the model as a subset of its support in which it is not possible to determine the cluster origin of a given sample with confidence higher than *∊*, that is

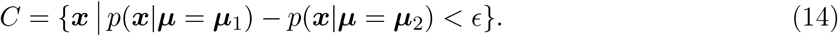

We can then consider clusters as connected if the “connected region” has probability mass greater than some threshold *α*:

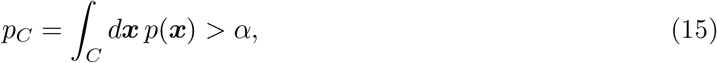

hence *p_C_* provides a measure of connectivity that measures how likely one observes “unassignable points” or “connecting points” when sampling from the cluster model. The corresponding empirical estimator for {***x***_*ι*_} and *n* observations

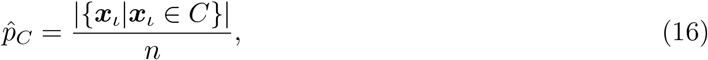

provides a measure for the connectivity of the empirical distribution, given the model assumption. Clearly, this is not yet a confidence measure for connectivity as arises from a hypothesis test. Testing the null hypothesis that *clusters are disconnected* requires to fix the parameter *δ*: then one can test whether the fraction of “connecting points” *p* ̂ _*C*_is significantly higher than *p_C_* as predicted the null model — if this is the case, one judges that the data is connected. However, fixing a parameter *δ* to some value can only be done in an ad-hoc way. Considering the alternative of testing the null hypothesis that *cluster centers are identical* does not provide an answer that uses the wished notion of connectivity. Finally, testing the null hypothesis that *clusters are connected* would require to specify a model class that models connectivity accurately, which would require a manifold-based model with continuous latent variables that parametrize the manifold. To circumvent these problems, we further investigate the estimator *p*̂ _*C*_as a direct measure for connectivity.

In particular, we want to compare *p*̂ _*C*_as arises from a cluster model with the subset of inter-cluster edges in the corresponding kNN graph. In the case of example (13), this amounts to determining the probability mass of the connected region

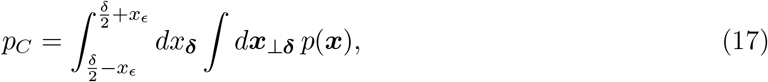

where points on the hyperplane ⊥ 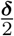 fulfill *p*(***x****|****μ*** = **0**) *− p*(***x****|****μ*** = ***δ***) = 0 and *x_∊_* determines how “thick” the “connected region” subspace around this hyperplane is. It is determined according to (14),

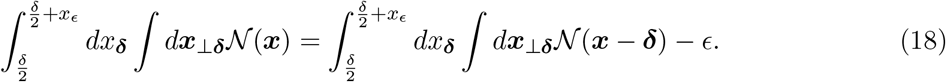

This can be easily solved by exploiting the radial symmetry of the integrand for the second term in ***x*** = *x*_***δ***_+ ***x***_⊥***δ***_in the *d –* 1 dimensional subspace. Using additionally the simple form of the isotropic Gaussian

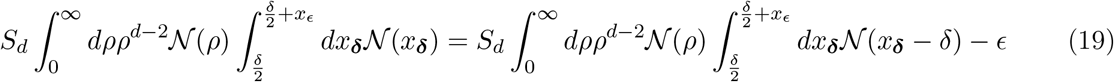

where 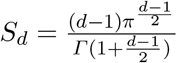 While this cannot be solved in closed form, the expression could be trivially further simplified using the error function erf. However, note the following.

While points that fall in the “connected region” surely qualify as inter-partition edges of the kNN graph that is partitioned according to the cluster labels, the assumption of a constant “thickness” *x_∊_* of the corresponding subspace is incompatible with kNN graphs. For large *δ* and large *ρ*, i.e. large distance of the “connected region” from the cluster centers, data points are more sparsely sampled and nearest neighbors are further separated. Hence, inter-cluster edges in a kNN graph correspond to different uncertainties *∊* about cluster membership depending on the value of *δ* and *ρ* and one has to assume *x_∊_ ≡ x_∊_* (*δ, ρ*). As a rough approximation, we estimate (17) using 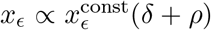 and obtain, assuming 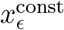 is small,

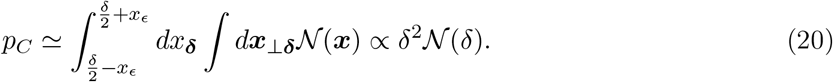

##### Benchmarks for Louvain-partitioned kNN graphs for clustering data

Let us study the result of connectivity measures (11) and (20) when applied to the Gaussian mixture model (13) in 20 dimensions. We consider 500 randomly generated datasets {***x***_*ι*_} of 100 observations ***x***_*ι*_. We consider both random partitions and Louvain partitions for increasing cluster distance *δ* in (13). The result is shown in Supplemental Figure 2:

a, Random partitions lead to the parabolic form of the number inter-edges predicted by (11) and the PAGA connectivity measure *c* is observed to vary between 0.5 and 1. The observed variance of *c* is the variance of the rescaled 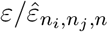 sum of Binomial variables, hence decreases as a square root for increasing values of *n_i_*.
b, Louvain partitioning a kNN graph deviates considerably from the random model already for cluster distance *δ* = 0, which is expected as the number of inter-edges is optimized to a local minimum. However, the PAGA connectivity measure *c* correctly corrects for the variation of *ε* with respect to partition sizes *n_i_*.
c, The summary statistics clearly shows that PAGA connectivity *c* is distributed with a much lower relative variance as compared to the number inter-edges *ε*.
d, Comparing the graph-based measures with the feature-space based measure of the frequency of points falling into a “connecting region”, one observes the expected qualitative agreement.

**Supplemental Figure 2.**
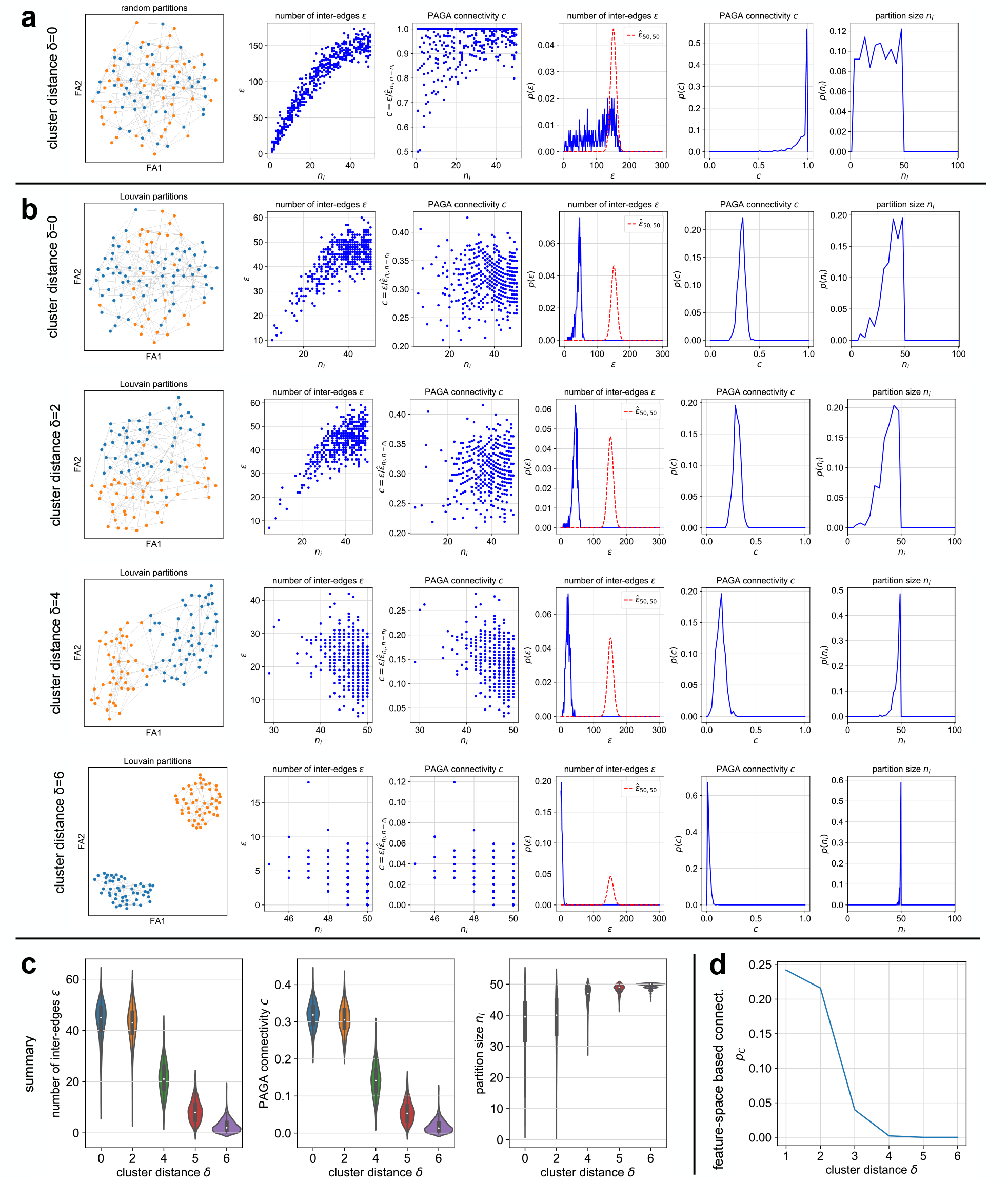
The PAGA connectivity measure provides a low-variance estimate of inter-partition connectivity that is independent of partition sizes. While in the first three rows, the number of inter-partition edges *ε* varies as a parabola with partition size *n_i_*, PAGA connectivity is independent *n_i_*. The figure is based on the Gaussian mixture model (13) and considers different cluster distances *δ*. From the model, we sample 100 observations ***x***_*ι*_in each of 500 simulations. **a,** Results for random bipartitions. **b,** Results for Louvain bipartitions. **c**, Summary of subpanels of **b**. **d,** Feature-space based estimate of connectivity (20).

**Supplemental Figure 3.**
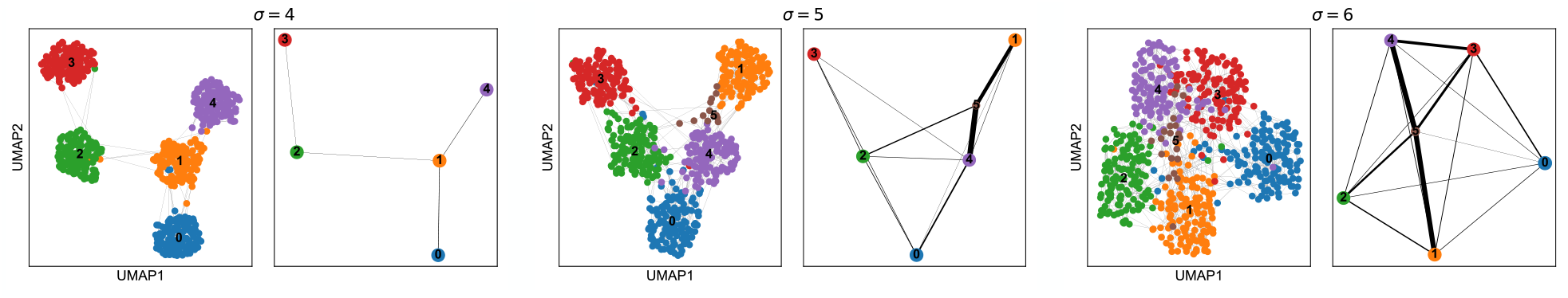
Connectivity of clusters sampled from Gaussian mixture model for different values of the standard deviation. Here, we show samples from three Gaussian mixture models, which display different degrees of clustering structure: the number of centers is fixed to 5 but the standard deviation *σ* is increases from 4 to 6. The figure makes evident that the same number of inter-edges for a small cluster leads to higher confidence in a connection than for a large cluster.

While Supplemental Figure 2c shows that the connectivity measure *c* of (11) has the desired property of showing low variance by virtue of the corrected partition-size effect, it would be desirable to have null model that correctly estimates the observed number of inter-edges of a Louvain-partitioned graph at cluster distance *δ* = 0. Evidently, such a model cannot be obtained as a simple expression but has to be fitted to data or obtained by sampling-based techniques. Independent of the computational burden introduced by this, which could hamper an efficient application in practice, there are many open conceptual problems of how to estimate the null model for real data, which is beyond the scope of this paper.

Let us finally inspect an example with several partitions sampled from a Gaussian mixture with five cluster centers. It can be seen that connectivity of clusters shows meaningful variation and reflects the basic assumption that a fixed number of inter-edges for small cluster gives higher confidence in its connection than for a large cluster (Supplemental Figure 3).

#### Supplemental Note 1.2: PAGA for mapping transitions between partitions

In this section, we briefly outline the generalization of the PAGA idea of abstracting from single-cell neighborhood relations to relations among groups by discarding insignificant relations attributed to a noisy graph.

In the context of RNA velocity [29], consider again a kNN graph in *d*-dimensional feature space {***x***_*ι*_} =*χ* = ℝ ^*d*^, given a distance measure *δ* such as Euclidean distance. Fitting a model for the steady state of reaction dynamics from unspliced to spliced RNA for each gene allows to define a velocity vector ***v***_*ι*_∈ ℝ^*d*^ for each cell *ι*. By computing the projection of the velocity vector onto the directions between the *k* neighbors of the cell in the kNN graph, it is possible to define a weight matrix *W* with entries

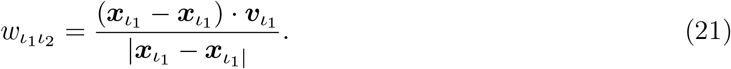

The resulting directed graph provides indication for that a cell transitions from node *ι*_1_ to *ι*_2_ with a transition tendency or strength 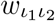. We note that *W* is not a stochastic matrix but simply the weighted adjacency matrix of a directed graph — hence we use the convention of adjacency matrices where 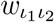 is associated with an edge pointing from *ι*_1_ to *ι*_2_. For a stochastic matrix, one usually follows the opposite convention (row vectors of probabilities and right stochastic matrices).

In order to judge whether a given group of cells *i* shows a significant drift or only random transitions to another group of cells *j*, we first need to correct for the group sizes *n_i_* and *n_j_*. Intuitively, one requires more transitions from *i* to *j* if *n_i_* is large. In order to correct for the size effect, we again consider the expected number of inter-edges from *i* to *j* under random sampling as in (7). This time, however, we do not consider the symmetrized version 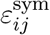 but consider *ε_ij_*. The estimator that is corrected for *n_i_* hence reads

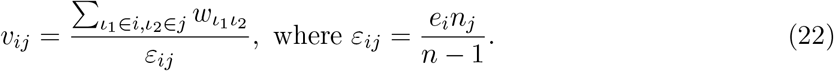

This is in complete analogy to (11) only that here, we consider edge weights that deviate from one. To judge whether the corrected summed difference of transition tendencies

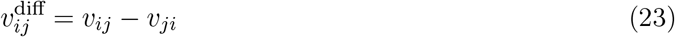

provide significant evidence for transitions to one group — deviates from 0 — we use a t-test. If 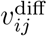 takes a positive significant value, we orient an arrow in the PAGA graph from *i* to *j*. The negative log p-value of the test has the desired property of taking large values if we can be confident in transitions from *i* to *j*. However, it approaches infinity for large values of 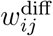, which is undesirable. Instead of the p-value, therefore, we use the summed transitions 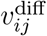 normalized to the standard deviation of the 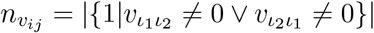 observations of

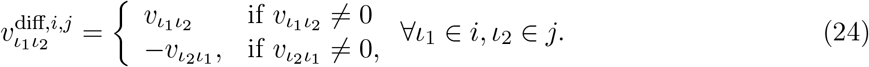

Hence, 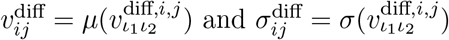 and we define the PAGA transition tendency as

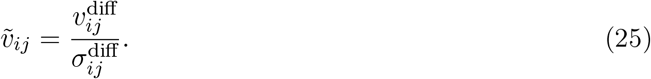

Examples are shown in Figure 3. We note that a completely different approach to modeling transitions between partitions has been proposed by David and Averbuch [37]. We also note that the approach here does not suffer from the problems discussed in [38]. Regarding the general interpretation of single-cell trajectories on snapshot data, we refer the reader to [8] and [38].

#### Supplemental Note 1.3: PAGA for multi-resolution analysis of data

Consider the finite node set *V* of observations of cells. To define a multi-scale graph, we assume an additional filtration {*V* ^(*i*)^}_*i*_on *V*, i.e. *V* ^(*i*)^ being a partition or clustering of *V*, with at its lowest level *i* = 0 being the whole set *V* ^(0)^ = {*V* }, and at its highest *i* = *n* being the set of nodes *V*^(*n*)^ = {{*v_ι_*}*| _ι_* ∈ *V* }, such that for *i >* 0

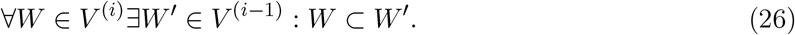

We interpret a low filtration level *i* as a coarse-grained clustering of observations, which means a low-resolution representation of the topology of data. Going to higher resolutions *i*, we aim to describe more fine-grained aspects of the data and eventually, for *i* = *n*, the single-cell level. Usually we are only interested in a small set of coarse-grained resolutions {*i*_1_*, i*_2_} that have a meaningful biological interpretation.

We can describe the filtration more explicitly by 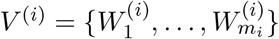, where by definition 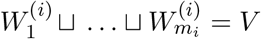 Then the partial order of the filtration induces a map

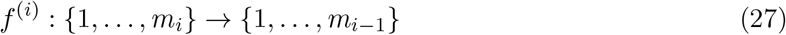

such that

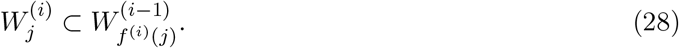

We can concatenate this to get for *i′ < i*

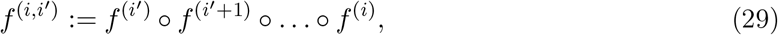

which lets us map any cluster on level *i* to a lower resolution *i′*.

**Supplemental Figure 4.**
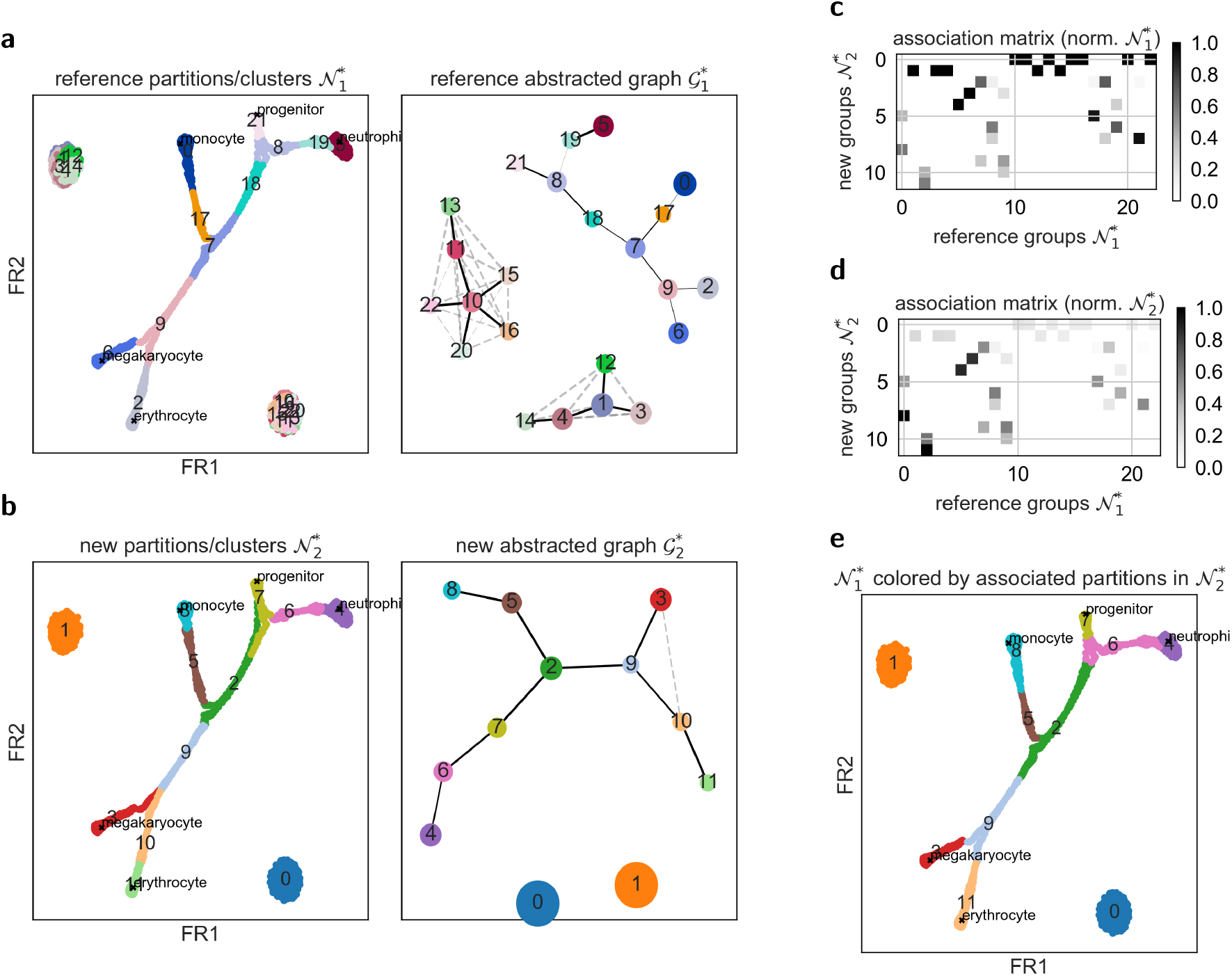
Illustration of multi-resolution analysis for simulated data. **a, b,** Partitions obtained using Louvain clustering in two runs with different parameters, equivalent to those shown in Figure 2a: both abstracted graphs describe the same topology. Note that in Figure 2a. c, d, Map between clusters of different resolutions. **e,** Reference partitions colored with the associated new partition that has the largest overlap.

Beyond Figure 3 and Figure 4, Supplementary Figure 4 shows a particularly simple example for a multi-resolution embedding for the simulated data of Figure 2. Supplemental Figure 4a and b show partitioned single-cell graphs and the associated PAGA graphs at two resolutions that differ from the one in Figure 2. Supplemental Figure 4c and d visualize the map (29) between clusters at different resolutions via association matrices. Supplemental Figure 4e shows the single-cell graph colored with mapped partitions.

Within PAGA, we combine this with an additional connectivity structure between vertices (*v*_1_*, v*_2_). We therefore assume a given graph *G* = (*V, E*) on *V* with edges *e* = {*v*_1_*, v*_2_} such that *v*_1_ ≠ *v*_2_ ∈ *V* . The edges may possibly be weighted with a function *w*(*e*) or be directed (*v*_1_*, v*_2_).

The filtration *V* ^(*i*)^ induces coarse-grained graphs *G*^(*i*)^ = (*V* ^(*i*)^, *E*^(*i*)^), where multiple definitions of coarse-grained edge sets could make sense. For instance, one could define the “complete abstracted graph” for resolution *i* by identifying *E*^(*n*)^ := *E* and demanding

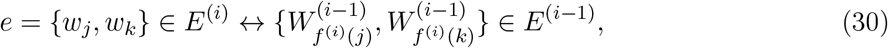

That is, any two supersets are connected if there is a connection one level below and hence on any level below. This is theoretically attractive since it is transitive, but is problematic in practice. It is not robust for noisy graphs and leads to strongly connected coarse-grained graphs that reflect the exact connectivity of the single-cell graph.

PAGA solves this by generating weighted coarse-grained graphs, which are then “abstracted” by thresholding low-weight edges. The simplest weight would be the corresponding number of inter-edges in the single-cell graph. Within PAGA, we use a statistical model to derive the weight (11) as the number of inter-edges in the single-cell graph divided by the expected number of inter-edges when assuming random connections (Supplemental Note 1.1).

Given an abstracted graph {*G*^(*i*)^} as defined above, we can now study paths across resolutions. We say that a low-resolution path 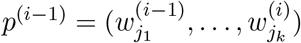 on *G*^(*i−*1)^ represents a high-resolution path *p^i^* on *G^i^* if

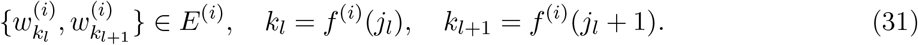

Conversely, we say that a high-resolution path 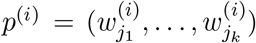 on *G*^(*i*)^ is represented by a low-resolution path *p*^(*i−*1)^ on *G*^(*i−*1)^ if

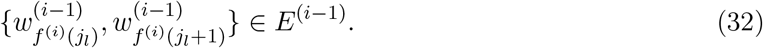

Note that the complete abstracted graph represents all high-resolution paths. However, a “good abstraction” of a high-resolution graph represents many high-resolution paths while being as sparse as possible. Sparsity arises naturally by demanding that the abstracted graph only represents those high-resolution paths that are statistically well supported, which is achieved through (11). This also defines the sense in which an abstracted graph can be said to be “topology preserving”.

#### Supplemental Note 1.4: Robustness of PAGA

Let us take a more practical view on the question of whether the topology of two abstracted graphs 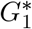 and 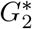agree under the constraint that the node labels of 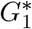 and 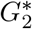 are consistent with each other. Moreover, instead of only detecting exact matches, we aim for a continuous measure of agreement.

##### Associating a partitioning with a reference partitioning

To establish such a measure, we first compute the overlaps of the partitions labelled by 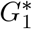 and by 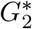 (Supplemental Figure 4a, b). By that, we generate non-unique associations between partitions, as visualized in an association matrix (Supplemental Figure 4c). The association matrix can either be normalized with respect to the reference groups 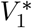 (Supplemental Figure 4c), with respect to the new groups 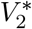 (Supplemental Figure 4d) or with respect to the union of partitions, which leads to the Jaccard index. Instead of the Jaccard index we want a score that measures how well two partitions mutually overlap — are mutually contained in each another — and consider the minimum of both mentioned normalizations — the “minimal overlap” — for each combination of groups 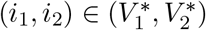. Supplemental Figure 4e colors each partition in 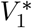 with the partition in 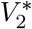 with which it has the largest minimal overlap.

##### Comparing paths in abstracted graphs

For each shortest path between two leaf nodes in 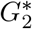, there is a shortest path between the associated nodes in 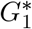 . This enables to compare the two paths and to count the fraction of steps that are consistent among two paths. To measure the agreement of the topologies between two abstracted graphs, we compute the fraction of agreeing steps and the fraction of agreeing paths over all combinations of leaf nodes in two given abstracted graphs.

For instance, consider the shortest path between leafs (21, 2) in the reference graph 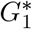 and the shortest path between leafs (7, 11) in the new graph 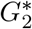 in Supplemental Figure 4a and b, respectively:

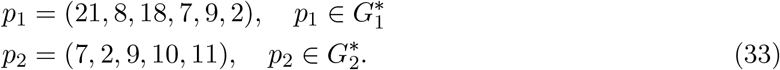

**Supplemental Figure 5.**
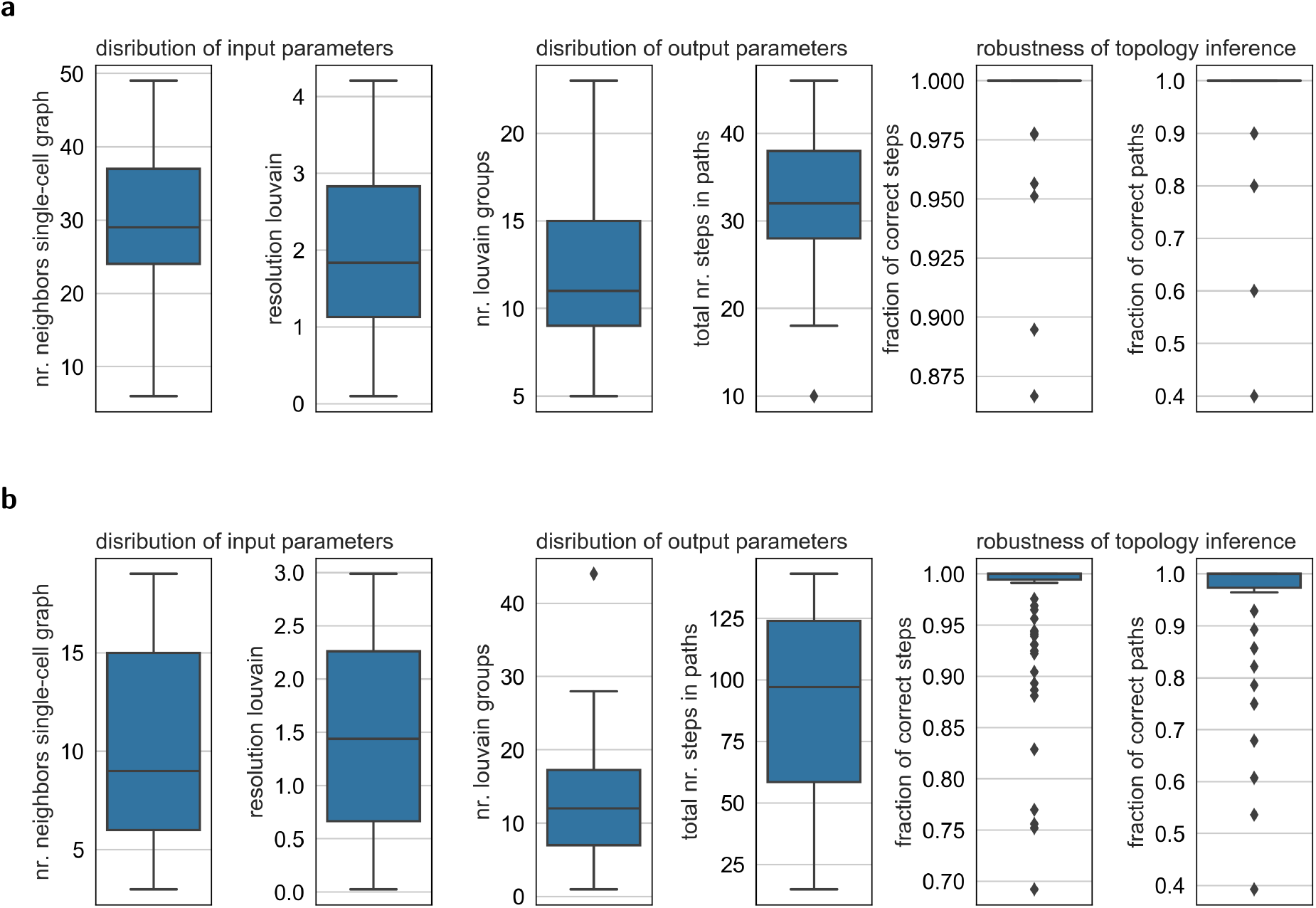
Robustness of PAGA. Sampling a wide variety of the input parameters (numbers of neighbors in the kNN graph and resolution of the Louvain partitioning) results in vastly varying numbers of partitions, hence vastly different clusterings of the data; note the large spread of the number of Louvain partitions. Nonetheless, the topology is robustly inferred. **a,** Simulated data as in Figure 2. **b,** Data of Reference [24] as in Figure 2.

By computing the overlap of reference partitions with new partitions, we can map *p*_1_ to the label space of 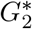

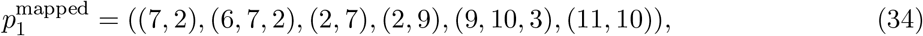

that is, partition 21 in *G*_1_ has finite minimal overlap with partitions 7 and 2 in *G*_2_, partition 8 in *G*_1_ has overlap with partitions 6, 7 and 2 in *G*_2_, and so on.

Transitioning through path *p*_2_ and counting for each transition whether it’s present or not in 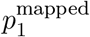 allows to count the number of agreeing steps. If all steps agree with each other, the paths *p*_1_ and *p*_2_ agree with each other. In the example of equation (33), *p*_2_ involves 4 steps, 4 of which agree with 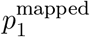.

##### Benchmark

In Supplemental Figure 5, we use the just-described measure to demonstrate robustness of PAGA.

##### A related measure from the literature

Previously, it has been suggested to correlate the distribution of path lengths of all paths through trees as a measure for topological similarity of trees [12]. Specifically, for a tree whose nodes label sets of data points, the lengths of all paths between all pairs of data points are computed. The correlation of such path-length sets obtained for two trees is suggested as a measure for topological similarity of the two trees. Besides being highly redundant and costly to compute, the resulting measure is very rough as it does not map paths onto each other; that is, it does not account for inconsistencies of paths with the same length.

#### Supplemental Note 1.5: Remarks on generating and partitioning single-cell graphs

At the heart of PAGA lies the assumption that the single-cell graph *G* — the kNN graph of observations ***x***_*ι*_in some feature space — provides a meaningful representation of data. This assumption is on one hand based on the community’s success with graph-based clustering [18–20], pseudotime inference [8], visualization [39, 40] and tSNE [16, 41]. On the other hand, it is based on the observation that neighborhood graphs robustly generalize any local distance measure to a global scale. As any fixed distance measure can at best encode a very rough notion of biological similarity with an exploding error for large distances, it is more robust to only evaluate it locally, and construct the global distances from the graph of neighborhood relations. See how some of us discuss this in more detail in Supplemntal Note 3 of Reference [15].

In this paper, we only consider established preprocessing steps [40, 42, 43] for single-cell transcriptomic data, each of which give rise to a different fixed distance measure. For single-cell imaging data. we consider a learned distance measure as induced by the feature space of a deep learning model [33]. Any other distance measure, for example, the kernel-based measure of Reference [44], or autoencoder representations [45, 46] would also be a viable option. Finally, we remark that denoising the kNN graph is another step, which should be considered. This can for instance be done by “pruning” [18] of by computing neighborhood relations in the truncated spectral approximation of the graph’s adjacency matrix (“diffusion map” representation).

A partitioning of *G* that maximizes the ratio of intra-to inter-partition edges is natural in the sense that it reveals regions of the graph with different connectivity and hence, different topology. Optimizing this ratio is known as optimizing modularity [35]. An efficient algorithm for this [19] has been suggested for single-cell biology by Levine *et al.* [18]. Loosely speaking, one expects to obtain the clearest coarse-grained group structure of data at a fixed resolution if choosing the partitioning that maximizes modularity. The original implementation of the Louvain algorithm could lead to disconnected communities when nodes were assigned to a common community with a single node connecting two or more parts of this community. When the central node was reconsidered by the algorithm and moved to a different community, a disconnected community remained. This unexpected behaviour could be fixed by splitting disconnected communities before each community aggregation step in the implementation of [47]. Note that the Louvain algorithm has also been adopted by popular single-cell analysis toolkits such as Seurat [42] and Cell Ranger [43]. Many other possibilities for partitioning *G* — or clustering the data — exist: we mention spectral clustering and the graph-based hierarchical clustering [48], which is based on a random-walk based distance measure.

#### Supplemental Note 1.6: Remarks on the reconciliation of clustering with trajectory inference algorithms

Here, we provide a more formal explanation of the discussion of Figure 1 in the main text. The aim of any pseudotime of given data is to provide a continuous latent variable that associates with continuous variation in the data; presumably the process that generated the data. Furthermore, pseudotemporal ordering of cells enables the identification of the relative timing of different events during the process — it tries to represent the internal “clock” of cells as encoded in its molecular configuration. A clustering analysis, by contrast, relates neither cells nor clusters to each other. With the PAGA graph *G^*^*, which describes the connectivity and “continuity” relations *c_ij_* of clusters *i* and *j* of *G*, and the pseudotime measure *d*(*ι*_1_*, ι*_2_), which measures the continuous progression of a cell *ι*_1_ to a cell *ι*_2_, one reconciles the result of a clustering analysis with the aim of a pseudotime analysis: Each cluster is related to any other cluster as either being disconnected or connected with one or several paths of high confidence in *G^*^*. Moreover, within each cluster, each cell is ordered according to pseudotime. One can hence trace a continuous process from a root cell *ι*_root_ in a root cluster *i*_root_ to any terminal cell *ι*_end_ in its terminal cluster *i*_end_ by following a path of high confidence (*i*_root_*, i*_1_*, i*_2_*, …, i*_end_) in *G^*^*. In each step of this path, the pseudotemporal ordering provides an ordering with single-cell resolution and, hence, one traces the progression of single cells along an ensemble of paths of high-confidence in *G*. Thereyby, PAGA provides a topology preserving map of cells as (*G^*^, d*). Without the PAGA graph *G^*^*, computing the ensemble of highly confident paths from from *i*_root_ to *ι*_end_ in *G* is a computationally much harder and an unsolved problem — only recently, during the revision of this paper, a simulation-based approximative approach has been proposed, but not validated on many datasets [49]. Presumably, the heuristics for their inference are less transparent and easy to control than the heuristics involved in partitioning a graph *G* and generating a PAGA *G^*^*.

### Supplemental Note 2: Random walks on graphs

On the single-cell level, the continuity of connections are believed to be well parametrized by a “pseudotime” [2, 3] that measures the distance covered in a continuous progression along a manifold. A robust kernel-based measure that can be easily extended to a graph, diffusion pseudotime, has recently been proposed by Haghverdi *et al.* [8]. This measure and similar scale-free random-walk based measures though do not account for clustering structure in the data; they are undefined for disconnected graphs. Below, we show how to overcome this limitation by extending these measures.

#### Interpreting random walks and their path distributions

It is important to note that in the whole paper, when we say “random walk on a graph”, we mean a discrete-space Markov process on the state space given by the nodes of the graph and non-zero transition probabilities between any two connected nodes.

Such random walks can be used to probe the global topology of the single-cell graph *G* but do not provide a good model for the biological processes that one might hypothesize to have generated the data in the first place. The primary deficiencies of the Markov random walk when seen as a model for a biological process are the following.

- Undirectedness. When progressing along a differentiation trajectory, at some point, one expects commitment of a cell to a specific fate and a directed motion to that fate with some fluctuations. By contrast, the diffusive motion induced by the Markov random walk is highly non-directed, which leads to unrealistic paths that go back and forth and pass through remote regions of the graph.
- Independence of the expression of specific genes. The random walk is independent of the expression of specific genes, which may be quite relevant for the commitment to specific fates; it only depends on global differences in the transcriptome.

These deficiencies of the random walk become apparent already when modeling a biological process using the simple stochastic differential equation based model discussed in Supplemental Note 5.3.

The distribution of single-cell paths that correspond to a path through the abstracted graph, by contrast, resolves the problem of undirectedness by bounding the distribution to the ribbon of the connected sequence of groups.

#### Existing random-walk based distance measures

For a single-cell graph *G* with *n*_nodes_ nodes and *n*_edges_ edges, consider the normalized graph laplacian [50, 51]

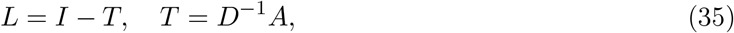

where *I* is the *n*_nodes_ × *n*_nodes_ identity matrix and *T* is the transition matrix of the same shape. *T* is obtained from the weighted adjacency matrix *A* of *G* by normalizing with row sums of *A*, that is, *D* is the diagonal matrix that stores the degree of each node in *G*. In practice, we compute the weights of the adjacency using a Gaussian decay with euclidian distance between two data points in gene expression space, see e.g. Reference [8]; after that, we density-normalize obtained weights [52, 53] as in Reference [8].

For a study of random walks generated by *T*, a spectral analysis of *L* and *T* is convenient and one hence considers the matrices *L ̃* and *T̃*, which are obtained by multiplying (35) with 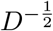 from the left and with 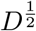 from the right

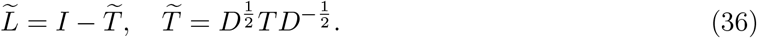

Hence, *L* and *L̃* have the same spectrum {1 *− λ*_1_, 1 *− λ*_2_*, …* } and the spectrum of *T* and *T̃* is given as {*λ*_1_*, λ*_2_*, …* } with *λ*_1_ = 1*, λ*_2_ *< λ*_1_*, …* for a connected graph *G* [50, 51]. For a disconnected graph with *n*_comps_ disconnected components, the adjacency matrix *A* has block-diagonal form with *n*_comps_ blocks and there are *n*_comps_ eigenvalues *λ_r_* with value 1 and corresponding eigenvectors that are the indicator vectors of the connected components. The eigenvectors *v* of *T* are related to the right eigenvectors *v ̃* of *T ̃* as [50–52]

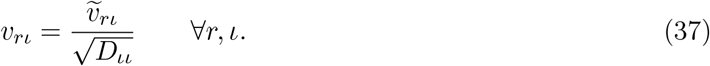

The right eigenvectors *v* of *T* are known as “diffusion map” coordinates [52], whereas the left eigen-vectors span the space of probability distributions of configurations of the Markov process. The first right eigenvector, corresponding to *λ* = 1, is the all-one vector — with only 1 as entry — and the first left eigenvector is the stationary state of the Markov process.

Using this notation, one obtains the mean commute time — the average number of steps one needs to arrive from node *ι*_1_ to another node *ι*_2_ — in equation (38a) [50]. One obtains “diffusion distance” [48, 52] in equation (38b) and “diffusion pseudotime” [8] in equation (38c).

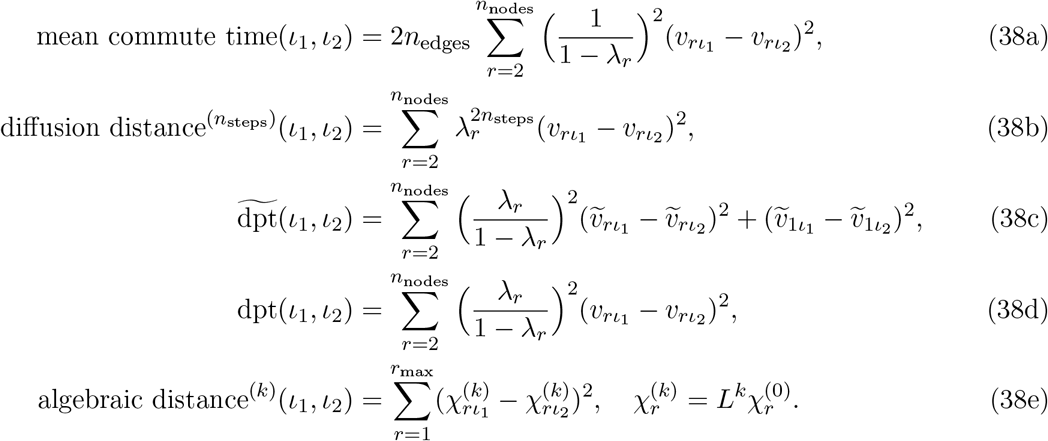

With equation (38d), we give a slightly altered definition of diffusion pseudotime, which is consistent with the other measures and was found to be equally-well performing for applications in single-cell biology by the authors of Reference [8] — note that using the *v_r_* basis instead of *v ̃_r_*, the last term in equation (38c) becomes zero. Highly related is algebraic distance on the graph as given in (38e) [see e.g. 53].

#### Interpretation of random-walk based distance measures

Random-walk based distances on graphs have first been used to cluster graphs in Reference [48] (38b) and Reference [54] (38a), albeit without considering neighborhood graphs of data points. Reference [52] proposed “diffusion distance” for measuring the similarity between data points, albeit not on a graph, but for a Gaussian kernel matrix. Then, a random-walk based distance measure for single-cell data has first been proposed to measure the similarity between cells by Reference [8]; again not formulated for graphs. These authors introduced the measure of equation (38c), which integrates out the number of steps *n*_steps_ in (38b) to arrive at a scale-free measure.

The dpt measure is highly similar to (38a), which is easier to interpret and scale-free, too: it measures the average number of steps it takes to walk from *ι*_1_ to *ι*_2_. While equation (38b) arises as the summed difference of transition probabilities to all other nodes for two random-walks of length *n*_steps_ that start at nodes *ι*_1_ and *ι*_2_, respectively [48, 52], (38d) considers the sum over all numbers of *n*_steps_, hence a difference of “accumulated transition probabilities”, which are difficult to interpret; the interpretation of equation (38c) is not easier.

Algebraic distance, which has been used for graph partitioning in recent years [53], approximates (38a) and diffusion pseudotime and provides the computationally most efficient way of computing a random-walk based distance measure.

#### Random-walk based distance measures for disconnected graphs

Evidently, both scale-free distance measures, mean commute time (38a) and diffusion pseudotime (38c), are not defined for a disconnected graph *G* for which *n*_comps_ *>* 1 eigenvalues are 1: they yield an infinite distance even for two nodes *ι*_1_ and *ι*_2_ that are in the same connected component of *G*. It is important to realize that each connected component of *G* automatically leads to a block *T_b_* in the transition matrix *T* that is itself a valid transition matrix and the spectrum of *T* is the union of the spectra of the blocks *T_b_*. The eigenvectors of *T* are the eigenvectors of the blocks *T_b_* filled with zeros at the positions of the other blocks [see e.g. 51]. Hence, we propose to extend mean commute time and diffusion pseudotime for disconnected graphs as

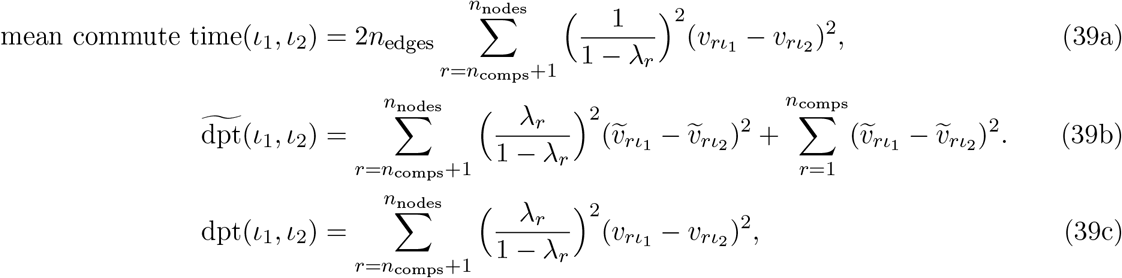

The distribution of zeros in the eigenvectors *v_r_* and *ṽ*_*r*_guarantees that for two nodes *ι*_1_ and *ι*_2_ in the same connected component *b*, only the spectrum of the block transition matrix *T_b_* contributes. For two nodes *ι*_1_ and *ι*_2_ in two disconnected components, the measures take the sum of their maximum values in both components, which should be interpreted as infinite. Without problem, one can make this explicit in the equations by distinguishing cases in which *ι*_1_ and *ι*_2_ belong to the same component from cases in which they belong to different components.

We note that, in practice, instead of summing over all eigenvectors *n*_nodes_, we sum over a low number of eigenvectors — “diffusion components” in the language of Coifman *et al.* [52] — as others [8, 54].

While in the present publication, we use equation (39b) throughout, we expect that equation (39a) could be useful in the future due to its easier interpretation.

### Supplemental Note 3: Comparisons with previous approaches

Establishing fair comparisons with other algorithms is difficult mainly for two reasons. (i) Comparisons on real data are problematic as a quantitative undebatable ground truth is hard to obtain. (ii) It is very easy to “make algorithms fail”, for example, by choosing an unsuitable preprocessing or pathologic parameters. Hence, after a comparison of concepts (Supplemental Note 3.1), we restrict ourselves to addressing fundamental problems and qualitatively wrong predictions of other algorithms for simple, simulated minimal examples with known ground truth (Supplemental Note 3.2), well-known real data, where we were able to reproduce published testing conditions [6] (Supplemental Note 3.3) and a comparison of runtimes (Supplemental Note 3.4). We note that a recent comprehensive trajectory inference review [4] positively mentions PAGA.

#### Supplemental Note 3.1: Conceptual comparisons

Monocle 2 [6] uses “reversed graph embedding” [55], which aims to fit a geometrical model for a graph to projections of the data to a low-dimensional latent space. Even though, in principle, any model could be used for that, in practice, only tree-like models are computationally tractable. Hence, Monocle 2 tries to force data into a tree-like topology without providing a statistical measure for how reliable the resulting fit is.

Spade [11], StemID 2 [13], Eclair [12], TSCAN [56] and Mpath [57] use different clustering algorithms such as k-means, k-medoids, hierarchical clustering or DBSCAN in a dimensionality-reduced space. In a second step, they fit a minimum spanning tree to either the centroid or medoid distances or to projections of cells on linear connections between centroids or medoids. In this, distances are computed using simple, fixed distance measures such as the euclidean or the correlation-based distance. Neither do these distances between clusters measure how well and if clusters are connected with each other, nor do these methods try to invoke a statistical model to address this question. The computationally expensive sampling procedures in StemID 2 and Eclair only partially alleviate the principle problem of high non-robustness that is caused by these deficiencies. Projections on linear connections between clusters assume a linear geometry of differentiation trajectories, which is certainly violated in practice. Hence, Mpath, for example, has only been shown to reconstruct processes with a single branching [57]. Moreover, it is important to note that none of the used clustering algorithms in these methods guarantees a topology preserving coarse-graining of the data: disconnected regions of data might cluster together and connected regions might be torn apart.

DPT [8] computes a random-walk based pseudotime for all cells. It cannot handle data with disconnected structure and is only able to detect single branchings which, in addition, is prone to violating the topological structure in the data (see, e.g., Figure 2c of Reference [10]). This problem becomes particularly pronounced in the extension of DPT to multiple branchings [34].

Rizvi *et al.* [10] suggest topological data analysis (TDA), in particular, the MAPPER algorithm [21] for analyzing single-cell data. MAPPER constructs a partial coordinatization of the data in the form of a simplicial complex, which has some similarity with the PAGA graph introduced in the present work. Both MAPPER’s simplicial complex and the PAGA graph represent connectivity of clusters in the data. However, the construction of MAPPER’s simplicial complex differs fundamentally from the PAGA graph. In particular, the clusters do not correspond to regions with controlled resolution and high intra-connectivity in the kNN graph, which are typically used in the field as proxies for cell types or cell states. Hence, in contrast to PAGA, MAPPER does not use an easily interpretable partitioning of the data into connected and disconnected regions, but a highly fine-grained, overlapping clustering, where clusters merely serve a technical purpose and are computed on a very low-dimensional map of data. Moreoever, MAPPER’s connectivity measure directly reflects the amount of overlap between these clusters. Hence, the measure does not induce a natural simplification by discarding statistically insignificant connections - it retains the full connectivity information of the overlapping clustering. Very generally, MAPPER is *not* based on simplifying a kNN graph of data points but uses the mentioned low-dimensional representation of data. Hence, MAPPER does not allow for a robust, random-walk-on-the-kNN-graph based distance measure for pseudotime estimation. Even though TDA allows the definition of continuous coordinates on the simplicial complex, their robustness and interpretability has not been shown. We interpret PAGA as a pragmatic, easily-interpretable, scalable and robust way of performing topological data analysis.

**Supplemental Figure 6.**
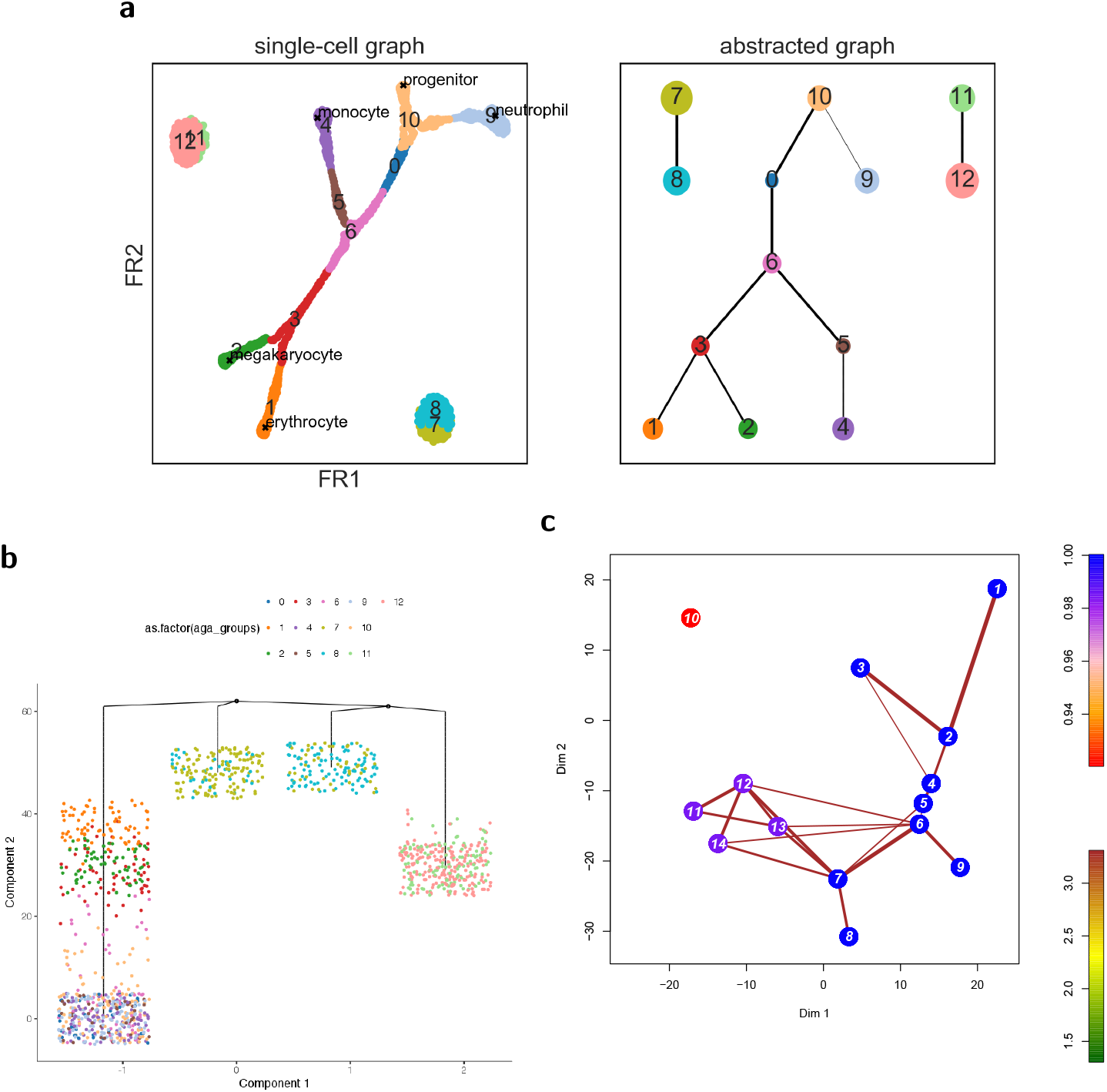
Comparison with Monocle 2 and stemID 2 for simulated myeloid differentiation and clusters. **a,** Prediction of graph abstraction, analogous to Figure 2. **b,** Prediction of Monocle 2 [6], the best result after testing several parameters for the latent-space dimension. The clusters (groups 7, 8 and 11, 12 in panel a) dictate the shape of the inferred tree, being responsible for three of the four observed branches. The continuous manifold is not resolved at all. The same coloring as in panel a is used. **c,** Prediction of the lineage tree of stemID 2, the successor of stemID [13]. The author of stemID, D. Grün, ran the simulation himself. The coloring and numbering of groups is chosen internally by stemID 2.

The graph coarsening approach of Wagner *et al.* [30] — developed at the same time and independently of PAGA — is also based on computing a connectivity measure based on the number of inter-edges between clusters in the single-cell graph. However, the approach has only been validated on a single dataset and does not provide a ready-to-use computational method for users. In addition, their metric computes for each pair of partitions the ratio of the number of inter-edges versus the number of the union of their out-going edges, which systematically overestimates connectivity. Assume two partitions share, relative to their size, a very small number of edges with each other and none with any other cluster - likely, these edges are a result of noise and the clusters are actually disconnected. However, in the approach of Wagner *et al.*, such clusters appear as strongly connected.

The hierarchical tSNE approach of Unen *et al.* [58] — published during the revision of the present paper — presents an idea for measuring “overlap between influence regions” of clusters obtained from density-based clustering. However, the measure is not related to the measure developed for PAGA. The authors proceed in using this overlap as a similarity measure to implement a hierarchical version of tSNE.

**Supplemental Figure 7.**
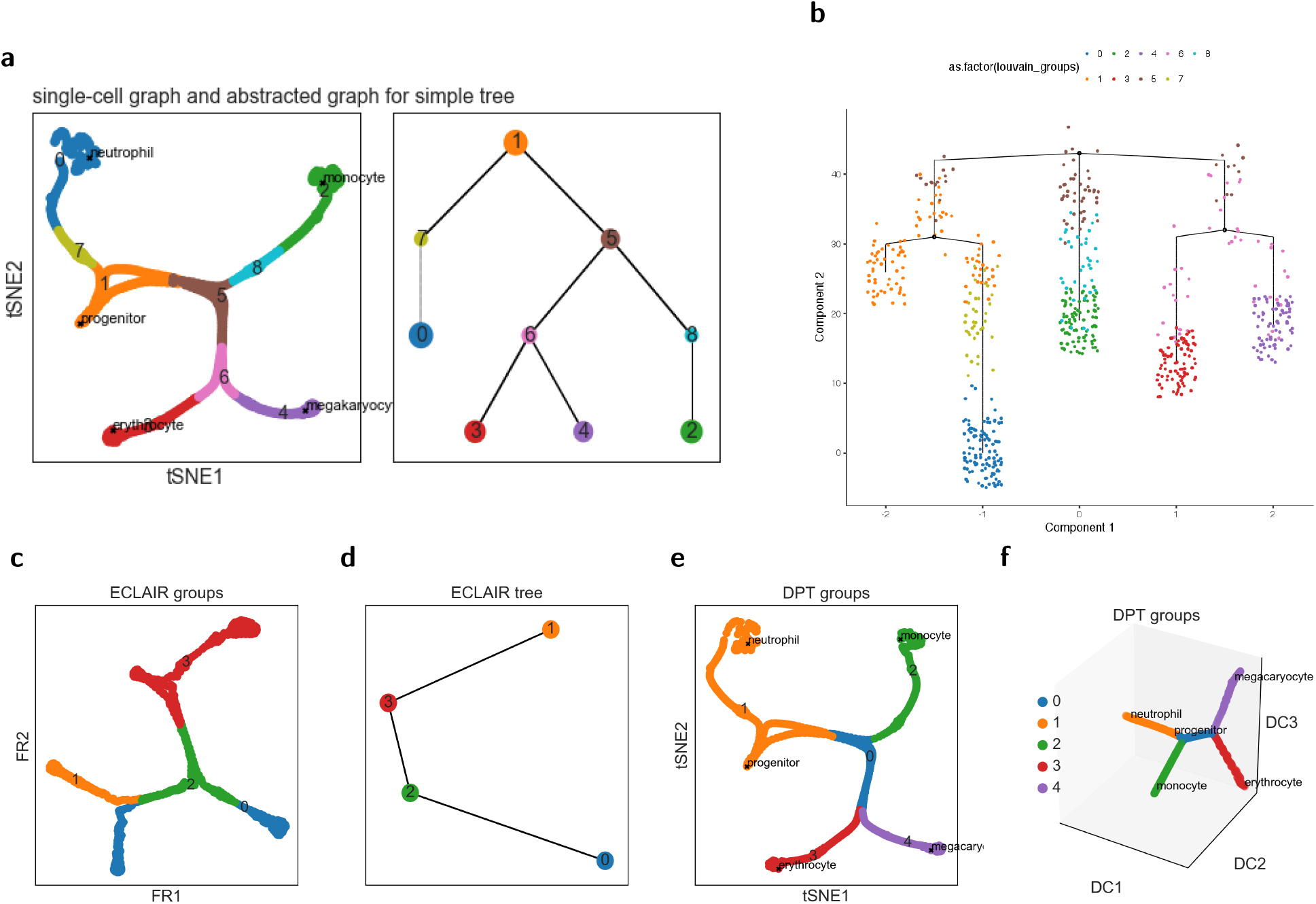
Comparisons for simulated myeloid differentiation giving rise to a simple tree-like manifold. Results using, **a,** graph abstraction, **b,** Monocle 2 [6], **c, d,** ECLAIR [12] and **e, f,** DPT [8] in its hierarchical implementation [34].

The graph-based approach p-Creode of Herring *et al.* [59] — published during the revision of the present paper — uses a density-adjusted kNN graph to produce an ensemble of potential trajectories. A consensus graph is then selected from the ensemble using a graph similarity metric.

We do not compare our method to Wishbone [7], which can only detect a single branching, nor to the fundamentally different, fully supervised approach STEMNET [60].

#### Supplemental Note 3.2: Simulated minimal examples with known ground truth

We consider a minimal example with known ground truth to show that graph abstraction overcomes qualitative conceptual problems in the design of algorithms for the inference of lineage trees. The dataset consists in a connected tree-like manifold and two disconnected clusters and has a clearly defined ground truth — a computational model for hematopoiesis (Supplemental Figure 11, Supplemental Note 5.3) — and very little noise. Nonetheless, none of Monocle 2, StemID 2, Eclair and DPT produce sensible results. Only when we removed the clusters from the data, these algorithms made sensible predictions. To reproduce the following comparisons and to get more information follow this link.

Graph abstraction recovers the ground truth (Supplemental Figure 6a). Monocle 2 [6] — even after testing several values for the latent-space dimension in Monocle 2 [6] — fits a tree to the clusters and misses to recognize the continuous manifold in the data (Supplemental Figure 6b). D. Grün ran stemID 2 — the unpublished successor of stemID [13] — on the minimal example. However, the produced graph-like object erroneously connects one of the clusters with the manifold (Supplemental Figure 6c). For the minimal example, we could not produce any sensible result neither with Eclair [12] — even after optimizing parameters in correspondence with the author G. Giecold — nor DPT [8].

**Supplemental Figure 8.**
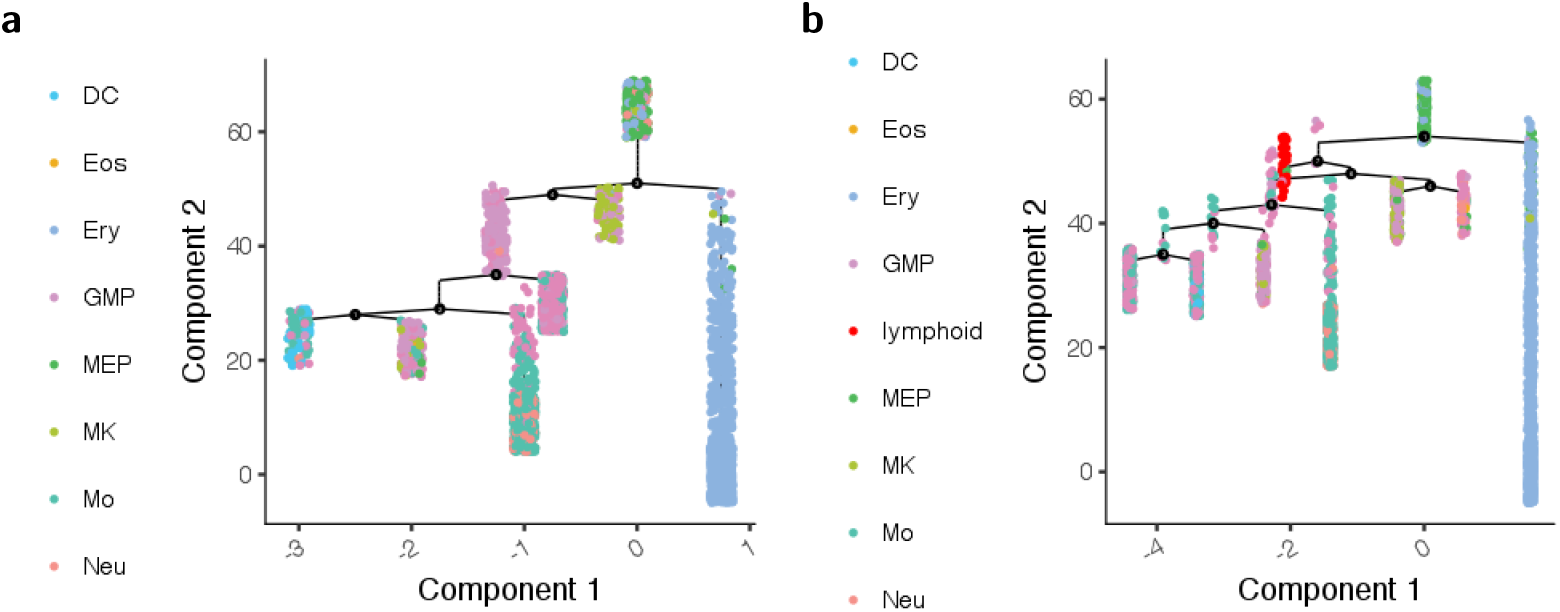
Monocle 2 for data of Paul *et al.* [24]. **a,** Monocle 2’s multiple branching example of Supplemental Figure 16 of Reference [6] using the same color coding as in the original publication. **b,** Rerunning Monocle 2 with the exact same parameters as for panel b, but keeping the lymphoids as in Figure 2a. The resulting tree changed dramatically and is no longer biologically meaningful. For example, the lymphoid cells are placed in the myeloid differentiation and myeloid progenitors (GMP) and monocytes (Mo) are distributed over all terminal states. As Monocle 2 does not provide confidence measures, the user erroneously expects all results to predicted with high confidence.

As a control, we aimed to obtain sensible results with the competing algorithms and considered a simpler dataset that only contains the continuous tree-like manifold of the previous example. Graph abstraction recovers the ground truth (Supplemental Figure 7a). Monocle 2 can be tuned — by adjusting the latent space dimension — to yield the correct result (Supplemental Figure 7b). Eclair [12] obtains a wrong result even for this simple tree (Supplemental Figure 7c, d). DPT [8] does, by construction, not infer a lineage tree but merely detects two branching subgroups; similar to a clustering algorithm. In a hierarchical implementation [34], it detects an arbitrary number of groups. Using the latter to detect four branchings we can detect two branchings (Supplemental Figure 7e) but fail to detect a third. Note that only when using diffusion maps for visualization, the clustering of groups appears natural (Supplemental Figure 7f).

#### Supplemental Note 3.3: Hematopoiesis

##### Comparisons for data of Paul et al. [24]

In the recent Monocle 2 paper of Qiu *et al.* [6] the data of Paul *et al.* [24] served as an example for the reconstruction of a complicated differentiation tree in Supplemental Figure 16. In the preprocessing step for the analysis of this data, Qiu *et al.* removed a cluster of lymphoid cells. In many situations, clusters of cells might not be annotated or not be clearly disconnected and it might not be clear whether one should remove them from the data. We therefore wondered what would happen when rerunning Monocle 2 with the exact same settings on the same data but keeping the cluster of lymphoids. While PAGA produces the same result irrespective of the presence of this cluster — it is simply disconnected in Figure 2, Monocle 2’s inferred tree changes dramatically and displays qualitatively wrong biology, for instance, by placing the lymphoid cluster in the center of the myeloid differentiation.

##### Comparison for data of Nestorowa et al. [25]

Supplemental Figure 9 shows a comparison for data of Reference [25].

**Supplemental Figure 9.**
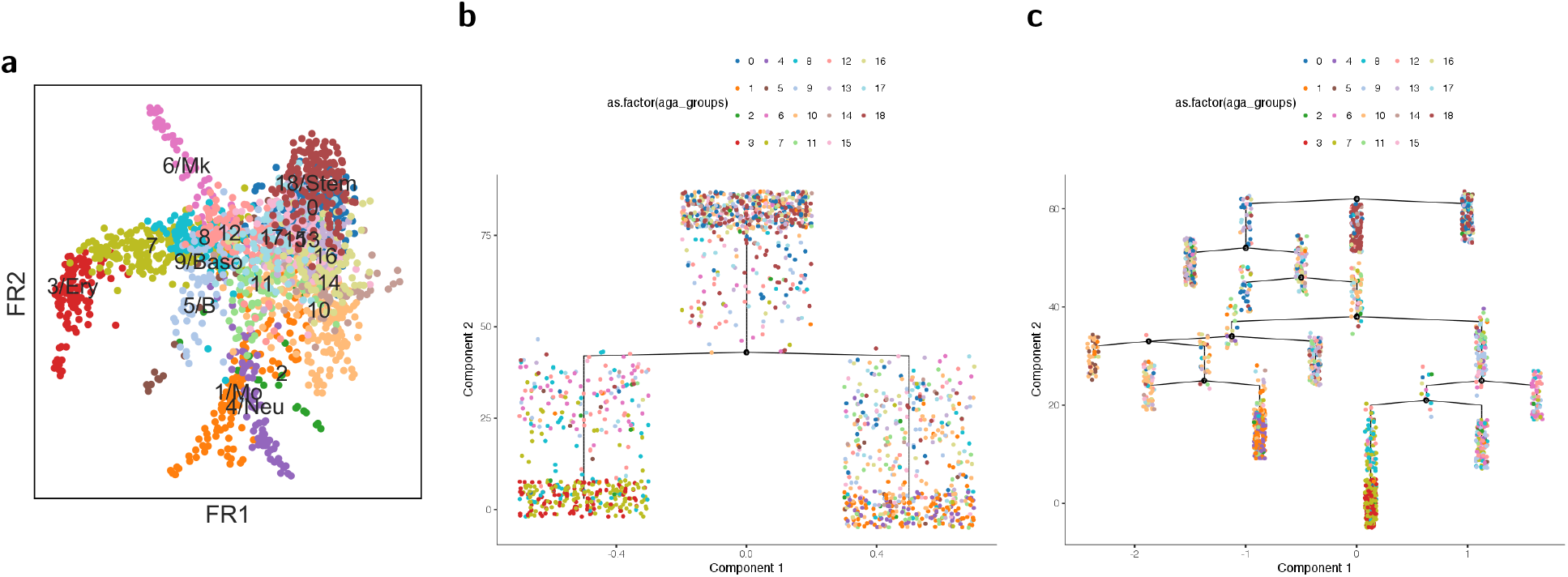
Comparison with Monocle 2 for data of Nestorowa *et al.* [25]. **a,** FR embedding colored by cell type annotation. This is an alternative embedding to the one shown in Figure 2 based on the graph that was not denoised. **b,** Running Monocle 2 for a latent space dimension of 4 underestimates the complexity of the differentiation manifold. **c,** Running Monocle 2 for a latent space dimension of 10 recovers the expected biology that late erythrocytes (3/Ery) and megakaryoctes (6/Mk) appear in the same region of the tree. Nonetheless, there are qualitative inconsistencies: neutrophils (4/Neu) and monocytes (1/Mo) appear in the same terminal branch. Megakaryocytes (6/Mk) appear in two branches. Basophils (9/Baso) appear as progenitors of erythrocytes and megakaryoctes and the disconnected B cells appear within the tree.

#### Supplemental Note 3.4: Runtimes

The authors of Monocle 2 report a runtime of 9 min for 8 000 cells [61] and a linear scaling. Extrapolation yields 76.5 min for 68 000 cells for which PAGA takes a few seconds — constructing the neighborhood graph and running clustering take an additional 3 min; hence PAGA is about 25 times faster.

PAGA for 1.3 million cells runs 90 s — constructing the neighborhood graph and running clustering takes about 45 min each. No other trajectory inference algorithm scales to such high cell numbers.

### Supplemental Note 4: Faithfulness of embeddings to global topology

Consider the cost function of the widely used tSNE algorithm [16],

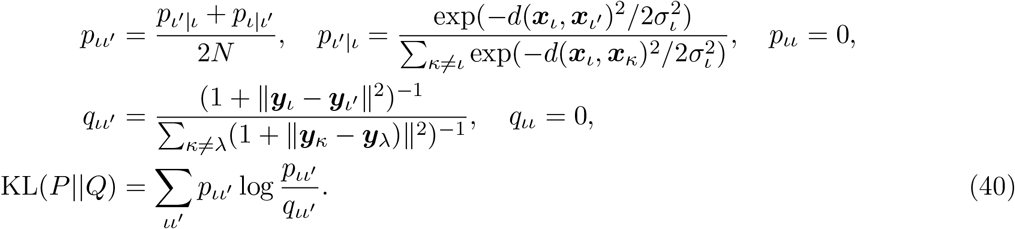

The double sum over *ι* and *ι′* is implemented as a sum over edges *e* = (*ι, ι′*) in the kNN graph of high-dimensional observations ***x***_*ι*_ ∈ *χ* . In the language of this paper, we say that *p_e_ ≡ p_ιι_′* quantifies the connectivity of node *ι* with *ι′* in the high-dimensional space *χ* and *q_e_ ≡ q_ιι_′* quantifies the connectivity in the embedding space 𝑌 . The optimized cost function hence is

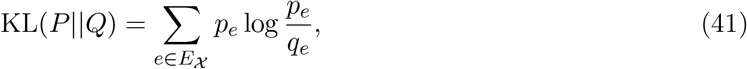

where we use the notation *E_χ_* to indicate that the edge set entering the optimization has been obtained as a kNN graph in *χ*.

#### Embedding cost functions as a binary classification problem

Let us take a different view on the quantification of how faithful the low-dimensional representation {***y***_*i*_} in 𝑌 is to the topology of the high-dimensional representation {***x***_*i*_} in *χ* . Let us define the ground-truth of this classification problem to be the kNN graph *G_χ_* = (*V, E_χ_*) fitted in *χ* . The state space of the classification problem is given by the edge set *E*_fc_ of the fully-connected graph *G*_fc_ = (*V, E*_fc_). We note that this is similar to the procedure introduced by [62].

In this classification setting, we require an embedding algorithm to predict for each edge *e* in *E*_fc_ whether it is an element of *E_χ_* . If it is an element, we assign the label *l_e_* = 1 to it, otherwise *l_e_* = 0. For each edge, the embedding algorithm makes prediction *l_e_* = 1 with probability *q_e_* and *l_e_* = 0 with 1 – *q_e_*. The standard cost function used to train such a classifier is the cross-entropy *H*(*P, Q*) or logloss, which is equivalent to the negative log-likelihood of the labels under the model

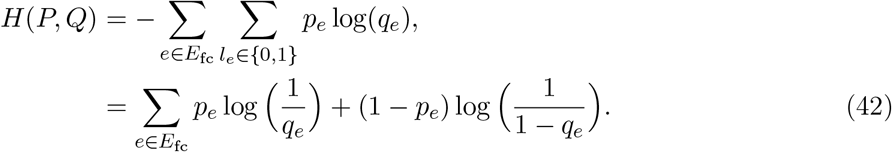

By virtue of 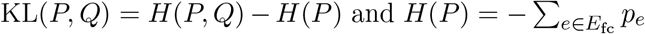, the KL divergence of predicted distribution *Q* and reference distribution *P* is

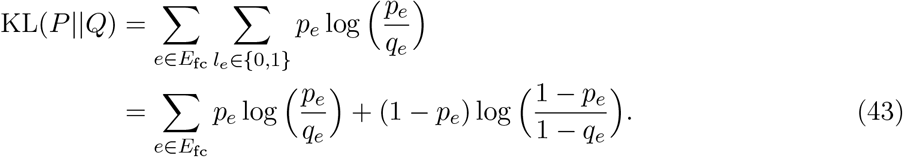

As in the optimization of the cost-function, the reference distribution *P* is fixed, it does not matter whether cross entropy or KL divergence is optimized.

Let us continue with interpreting the KL divergence, which both appears in UMAP and tSNE — however, in the case of tSNE the second term in (43) is absent. We can interpret the two terms in the KL divergence as follows

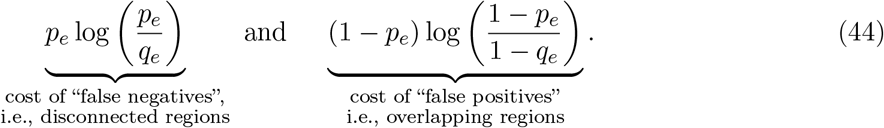

The first term generates cost that can be attributed to false negatives: edges in *E_χ_* that are not “detected” by the embedding algorithm and hence miss in the predicted edge set *E_𝑌_* . This occurs when a pair of points (*ι, ι′*) is far apart in the embedding space and hence has low or zero predicted connectivity *q_ιι_′* but has high connectivity *p_ιι_′* in the reference space *χ* . As the cost function diverges if any *q_e_* = 0 if the corresponding *p_e_* ≠ 0, such disconnected structure should in theory not occur. However, it is well-known that tSNE produces many spurious disconnected structures in the embedding — this can be attributed to the fact that values for *q_e_* have to be clipped to finite values so that the cost function itself remains finite and can be numerically stably optimized. Also UMAP [22] suffers from this problem.

The second term in (43) can be attributed to false positives: edges predicted by the embedding algorithm even though they miss in *E_χ_* . This occurs in “overlapping regions” of the embedding, where *q_e_* is close to 1 even though *p_e_* is close to 0. This phenomenon is frequently encountered in graph drawing algorithms such as ForceAtlas2 [23].

#### A novel cost function that accounts for global topology

Both disconnected and overlapping structure in the embedding present strong violations of the global topology represented by *G_χ_* that hinder interpretability by humans. However, in the cost function (43), these violations only contribute as strongly as violations of local topology with a weight of order

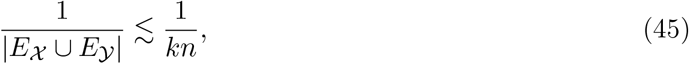

where the right-hand-side estimate holds for kNN graphs. Hence, for high numbers of observations *n*, the cost of violating global topology approaches zero, even though this is in stark contrast to what is desirable to the human interpreter.

**Supplemental Figure 10.**
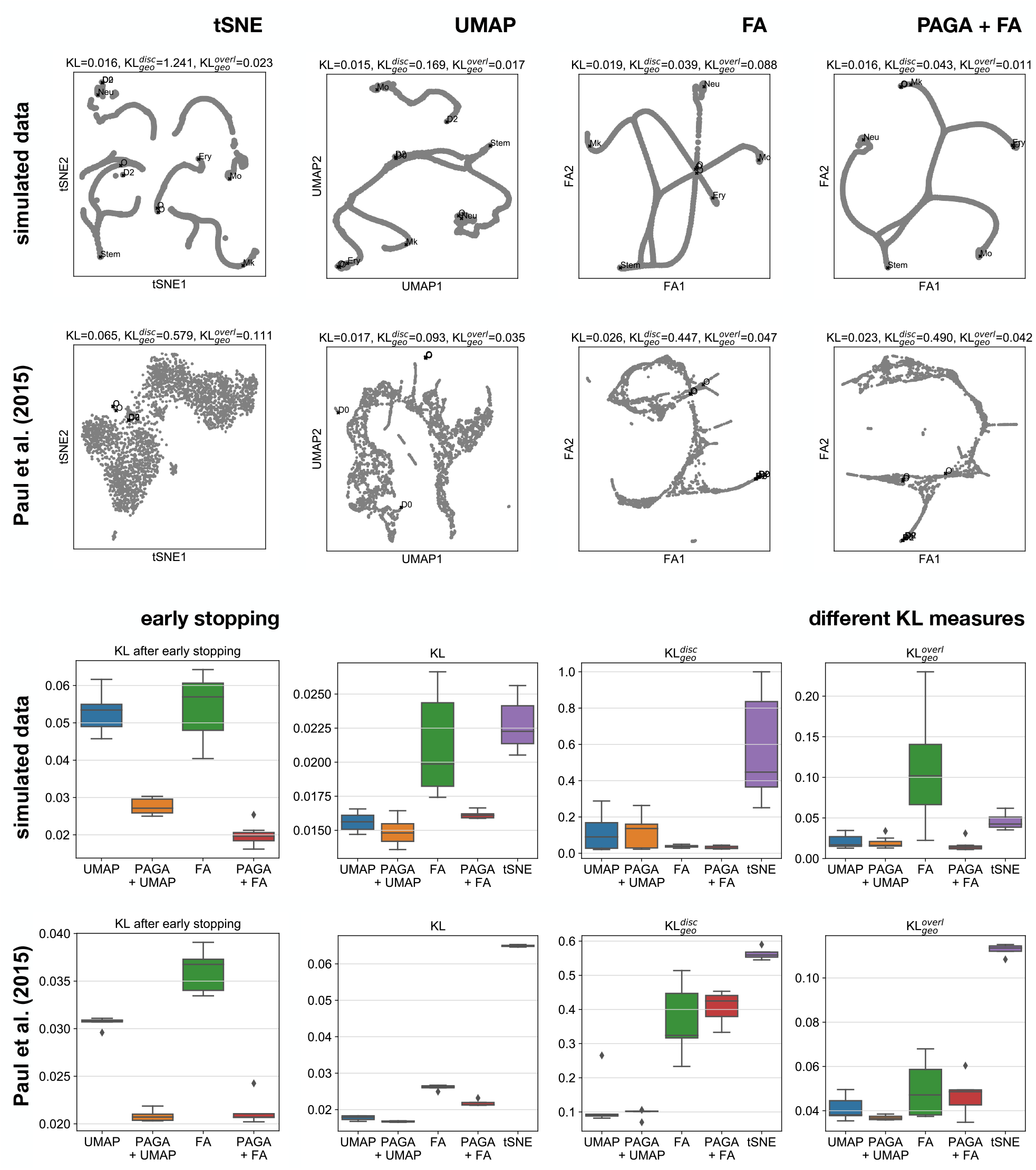
Performance of different embedding algorithms in representing of local and global topology of high-dimensional data. Using PAGA as an initialization for established manifold learning algorithms both leads to the best quality embeddings and enables early stopping. See (43) for the definition of the conventional KL divergence, which quantifies the preservation of local topology, and (46) for geodesic KL divergence KL_geo_, which also accounts for preservation of global topology. Highlighted are points in the embedding that lead to strong violations of global topology (overlapping “O” and disconnected “D” points).

In order to remedy this discrepancy, we suggest a weighted KL (or cross-entropy), which reflects the desire that edges that violate the global topology carry a higher weight than edges that violate local topology. Specifically, we suggest

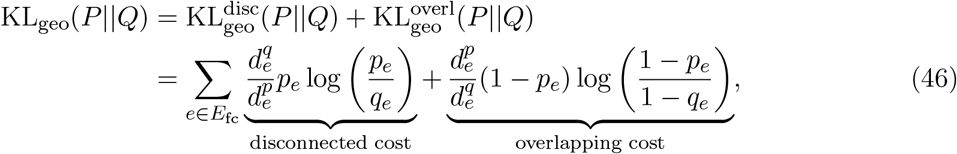

where 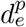 and 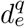 denote random-walk based distances in the kNN graphs *Gχ* and *G 𝑌* and are hence estimators of geodesic distances of the manifolds *χ* in and *𝑌*. See an extensive review of such distances in Supplemental Section 2.

Clearly, geodesic distance captures important aspects of the global topology of a manifold. The interpretation of the factors 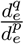 and 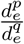 is hence as follows. If there is a globally disconnected region in the embedding, this causes 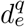 to diverge to infinity. If the region is also disconnected in the high-dimensional reference space, the effect cancels out in 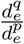, otherwise, the violation receives a high weight in (43). The argument is analogous for overlapping regions.

#### PAGA provides faster convergence and more interpretable single-cell embeddings

Throughout this paper, established manifold learning algorithms only provided embeddings that would violate the topology of data found in the high-dimensional feature space, see for instance Figure 3. Using (46), we can now quantify these violations and show that PAGA-initialized manifold learning both provides embeddings that are more faithful to the global topology and allows faster convergence also with respect to the conventional cost function (43). The results are summarized in Supplemental Figure 10) for the first two examples shown in Figure 2.

1. The two rows showing different embeddings highlight globally disconnected points (marked as D0, D1, …) and globally overlapping points (marked as O0, O1, …) identified by maximizing the weight factors 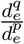 and 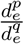, respectively. The title of the embeddings show the conventional KL divergence and the reweighted geodesic KL_geo_ introduced in (46). Quantitative values agree with the visual impression except for the FA embedding of Paul *et al.*, which we would expect to have a higher 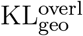.
2. The two rows showing statistics of KL measures for different embeddings and with or without initialization with PAGA have been produced by rerunning embedding algorithms 10 times. They show that either PAGA+FA or PAGA+UMAP achieve the best values throughout. The right-most panel shows KL values after early stopping, illustrate that already after 50 optimization epochs, KL values comparable to the converged result (*≥*200 epochs) are obtained.

### Supplemental Note 5: Datasets

#### Supplemental Note 5.1: Simulated dataset for hematopoiesis

We use a literature-curated qualitative – boolean – gene regulatory network of 11 genes that aims to describe myeloid differentiation [64] and has been used for benchmarking the reconstruction of gene regulatory network from a single-cell graph of state transitions in Reference [65]. The boolean network evolves according to

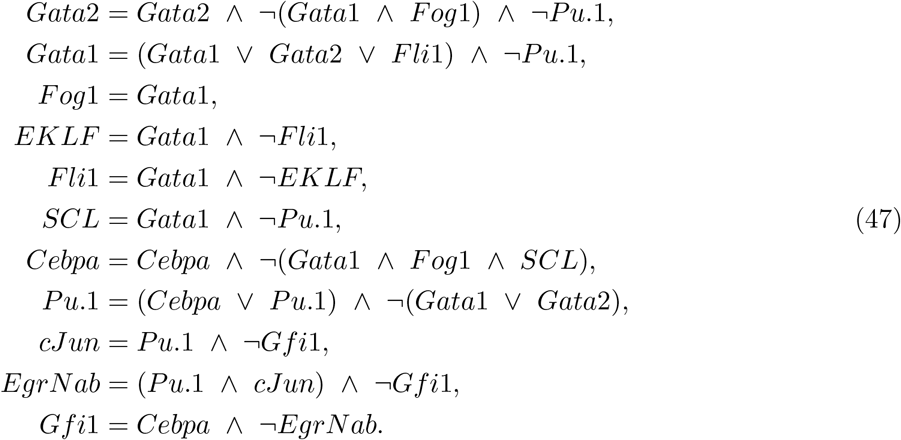

**Supplemental Figure 11.**
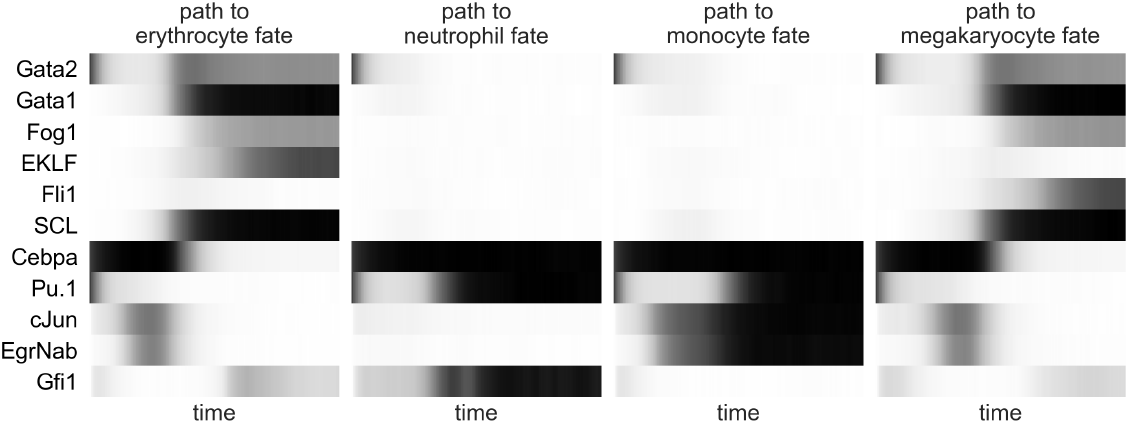
Simulated data for myeloid differentiation. Four representative realizations of time series representing differentiation to four different fates. These time series have been sampled from (47) after transformation to a system of stochastic differential equations Reference [34, 63].

These boolean equations are translated into ordinary differential equations following Reference [63]. Within Scanpy [34], they are simulated as stochastic differential equations by adding Gaussian noise.

Simulations result in four classes of realizations of gene expression time series, each of which corresponds to the convergence to an attractor that represents a certain cell fate of myeloid progenitors: erythrocyte, neutrophil, monocyte and megakaryocyte (Supplemtal Figure 11). We concatenate four typical realizations (Supplemtal Figure 11c, d) with 160 time steps, which yields 640 data points in total.

To model clustering, we sample 640 data points from a Gaussian mixture model with two Gaussians and random centers in an 11-dimensional space. The minimal dataset of Figure 2 and Supplemental Figure 11 consists of the concatenated data matrices of the simulated myeloid progenitor development data and the Gaussian mixture model, corresponding to 1280 cells.

#### Supplemental Note 5.2: One million neurons

As an input for the PAGA analysis, we used the kNN graph obtained by running the *pp.recipe_zheng17* [43] preprocessing function within Scanpy [34] and computing neighbors on 50 principal components. See Supplemental Figure 12 for visualizations of these data using PAGA and UMAP.

#### Supplemental Note 5.3: Experimental datasets for hematopoiesis

For preprocessing the data of Paul *et al.* [24], we used Scanpy’s preprocessing function *pp.recipe_zheng17*, for the data of Nestorowa *et al.* [25], we used *pp.recipe_weinreb17*. For the data of Dahlin *et al.* [26], we used the preprocessing of the original publication. We then computed kNN graphs on 20 principal components with *k* = 4 for Paul *et al.* and Nestorowa *et al.* and for *k* = 7 for Dahlin *et al.*, as in the original publication. PAGA can be applied on the resulting kNN graphs and yields meaningful results. However, for Figure 2, we further denoised the graph by approximating its adjacency matrix with the first 15 spectral components. We performed this approximation by recomputing a kNN graph using the first 15 diffusion components of the PCA-based graph. For this recomputation of the kNN graph, we used *k* = 10 for Paul *et al.* and Nestorowa *et al.* and for *k* = 15 for Dahlin *et al.*. We note that denoising the kNN graph by a different technique has already been suggested in Reference [18].

**Supplemental Figure 12.**
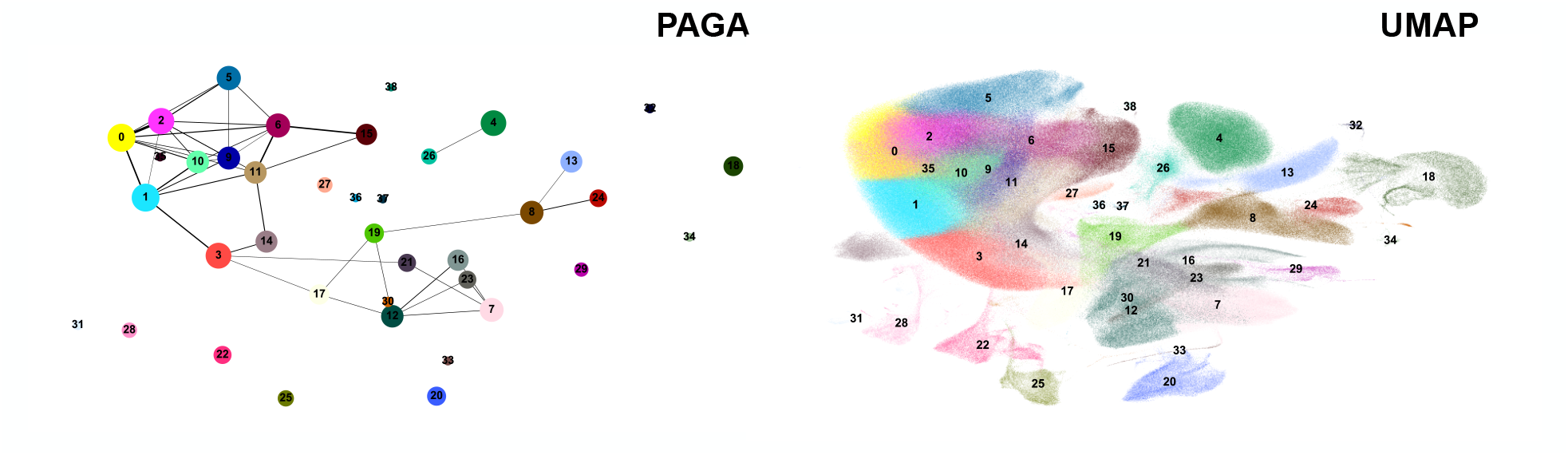
PAGA and UMAP on 1.3 million neurons dataset from 10x Genomics [31]. While PAGA takes about 90 s of computation time, UMAP takes about 3 h. Due to overlaying groups, UMAP blurs the topological structure and visually suggests too much connectivity — it suggests connections where there actually are none, as shown by PAGA. Consider the example of cluster 19, which UMAP suggests to connect to cluster 21 whereas it actually connects to clusters 17, 30 and 8. Note that this figure is the only instance of the main text in which we used the default initialization of UMAP and not the PAGA coordinates.

**Supplemental Figure 13.**
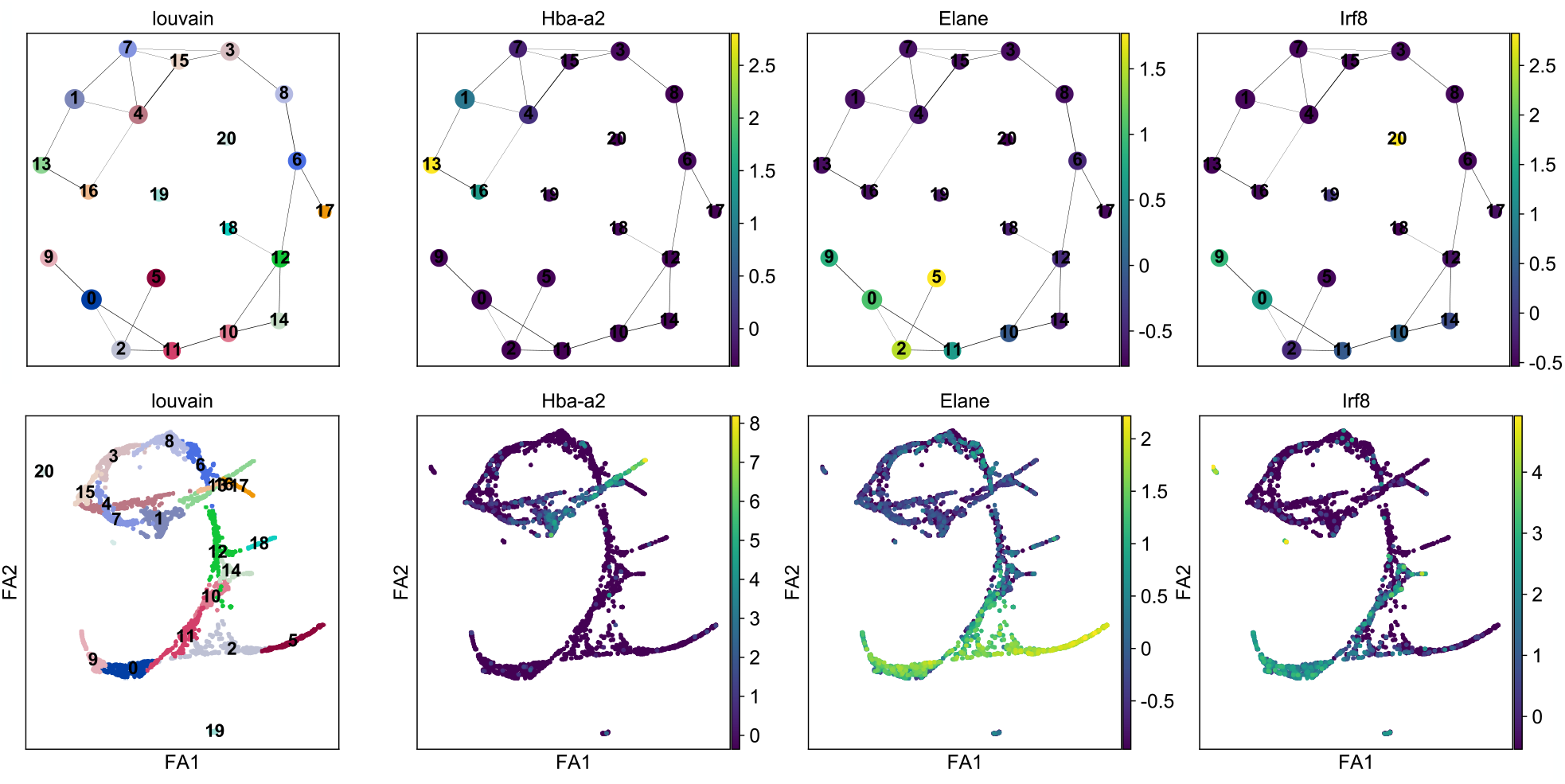
Annotation of Louvain clusters for hematopoietic data of Paul *et al.* using PAGA and ForceAtlas2. PAGA and the ForceAtlas2 (FA) embedding were computed with default parameters. In contrast to the PAGA-initialized embedding of Figure 2, the single-cell layout shows overlapping structure. While this is a relatively small and simple dataset an interpretable single-cell embedding, it exemplifies that the PAGA graph can be used as an even easier accessible visualization of the data. Both PAGA and single-cell graph show the Louvain clusters, an erythroid branch marked by *Hba-a2*, a neutrophil branch marked by *Elane* and a monocyte branch marked by *Irf8*.

**Supplemental Figure 14.**
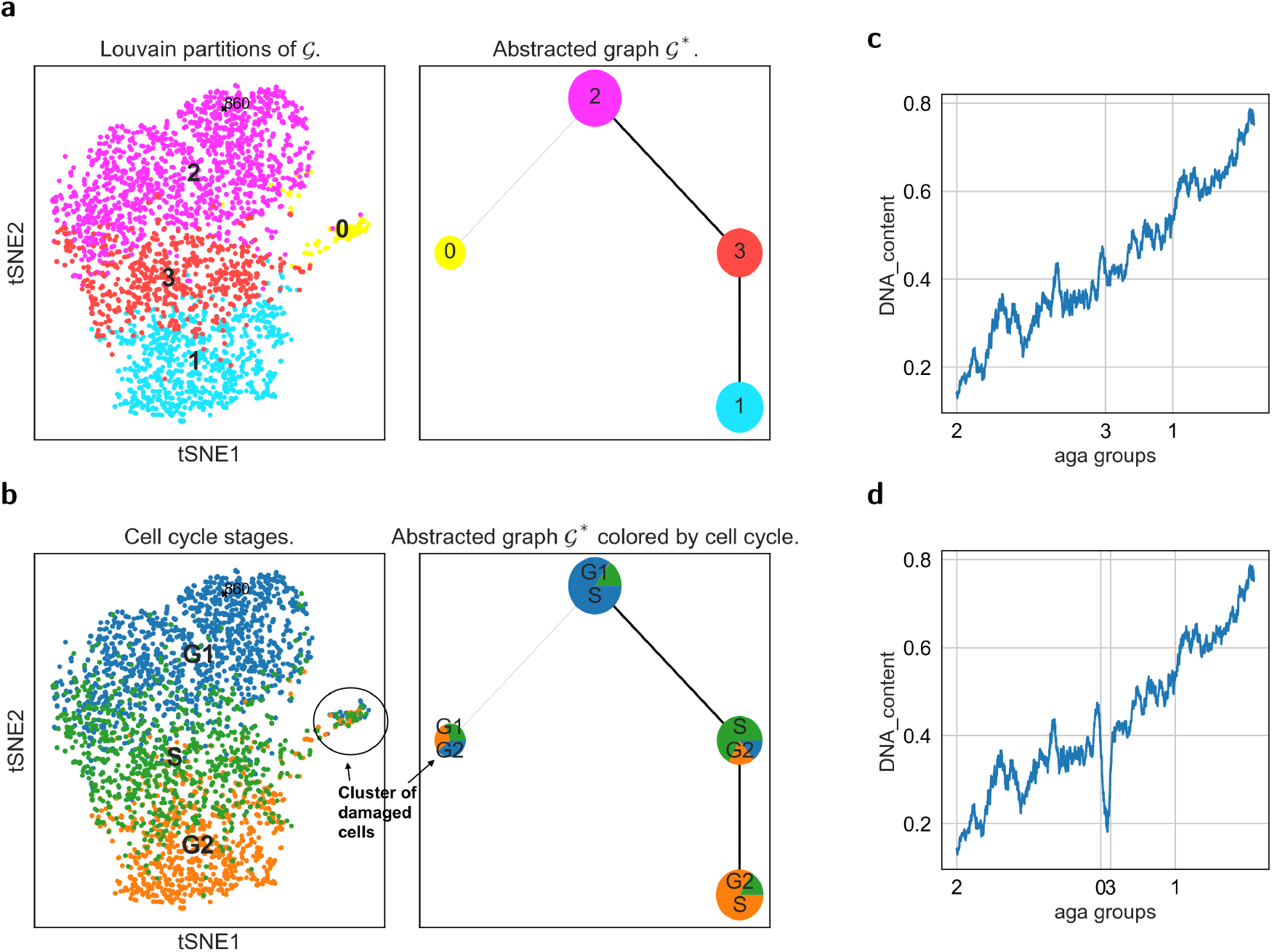
Abstracted graph for deep learning based feature space. Analyzing single-cell images via a deep learning based distance metric. Graph abstraction correctly recognizes the cluster of damaged cells as not belonging to the biological path that corresponds to cell cycle evolution through the interphases G1, S and G2. **a,** Abstracted graph with Louvain partitions. **b,** Associated cell cycle phases. **c,** DNA content along a valid path in the abstracted graph. **d,** The DNA content along an invalid path that involves the damaged cells shows a clear non-biological kink.

See Supplemental Figure 13 for an example of annotating clusters using PAGA for Paul *et al.* [24].

#### Supplemental Note 5.4: Planaria

For the analysis of the Planaria data of [15], we used the preprocessing of these authors. We computed a kNN graph on 30 principle components with 30 neighbors.

#### Supplemental Note 5.5: Zebrafish embryo

As an input for the PAGA analysis, we used the kNN graph and clustering of Wagner *et al.* [30] as provided by the authors.

#### Supplemental Note 5.6: Deep-learning-processed image data

Without extensive preprocessing, the graph of neighborhood relations of data points in gene expression space is useless if computed with a simple fixed distance metric (euclidian, cosine, correlation-based, etc.). If one considers the pixel space of images the problem is even worse and it is impossible to come up with preprocessing methods that lead to a meaningful distance metric. It has recently been shown that a deep learning model can generate a feature space in which distances reflect the continuous progression of cell cycle and a disease [33], that is, deep learning can generate a feature space in which data points are positioned according to biological similarity and by that generates a distance metric that is much more valuable than a simple fixed distance metric. We demonstrate that graph abstraction is useful for reconstructing the cell cycle from image data while and identifying a cluster of damaged cells (Supplementary Figure 14).

## Footnotes

1 This is a simple expression in the bipartitioned case *h* = *h_i_* + *h_j_* 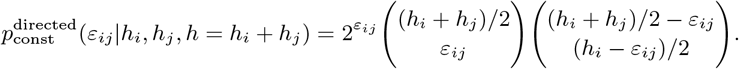

